# TMEM209/NPP-28 partitions PLK-1 between the nuclear envelope and nuclear interior during mitosis

**DOI:** 10.64898/2026.10.05.752757

**Authors:** Victória Puccini de Castro, Sarah R. Barger, Razvan Azamfirei, Tiffany Y. Su, Rebecca A. Green, Ronald J. Biggs, Frances E. Q. Moore, Shoken Lee, Karen Oegema, Pablo Lara-Gonzalez, Shirin Bahmanyar

## Abstract

Nuclear envelope breakdown (NEBD) requires disassembly of nuclear pore complexes (NPCs). Polo-like kinase 1 (Plk1/PLK-1) binds and phosphorylates NPCs to initiate NEBD. Plk1/PLK-1 also accumulates in the nucleus to regulate chromosome segregation. How Plk1/PLK-1 is spatially partitioned between NPCs and the nuclear interior is unknown. Here, we show NPP-28, the *C. elegans* homolog of transmembrane nucleoporin TMEM209, docks PLK-1 at the nuclear envelope and limits its nucleoplasmic accumulation. Deletion of *npp-28* or mutation of PLK-1 docking sites reduces nuclear envelope-associated PLK-1 and increases its nucleoplasmic levels. Accordingly, *npp-28* loss delays NEBD and promotes anaphase onset. PLK-1 phosphorylates NPP-28 *in vitro,* yet mutation of predicted PLK-1 phospo-sites in NPP-28 accelerates NEBD. Thus, unlike other nucleoporin substrates of PLK-1, NPP-28 phosphorylation prolongs NEBD. We also find context-dependent functional relationships between NPP-28 and other transmembrane nucleoporins. NPP-28 thus partitions PLK-1 between the nuclear envelope and nuclear interior to coordinate NEBD with anaphase onset.

## Introduction

Animal cell mitosis begins with nuclear envelope breakdown (NEBD). NEBD requires disassembly of nuclear pore complexes (NPCs) and the nuclear lamina. The mitotic kinase Polo-like kinase 1 (Plk1/PLK-1) binds and phosphorylates nucleoporins to dismantle NPCs (Kutay et al., 2021; Laurell et al., 2011; Linder et al., 2017; Martino et al., 2017; Nkombo Nkoula et al., 2023; Velez-Aguilera et al., 2020). Plk1/PLK-1 binds nucleoporins through its polo-box binding (PBD) domain and phosphorylates specific linker nucleoporins to disrupt their association with the NPC scaffold (Laurell et al., 2011; Linder et al., 2017; Martino et al., 2017; Nkombo Nkoula et al., 2023). Plk/PLK-1 also accumulates in the nucleus to regulate chromosome segregation from kinetochores. How Plk/PLK-1 pools are distributed between NPCs and the nucleoplasm is not understood.

Transmembrane nucleoporins anchor NPCs to the pore membrane (Antonin et al., 2005; Onischenko et al., 2009; Stavru et al., 2006b, 2006a). In human cells, Plk1 disrupts membrane anchoring of NPCs during mitosis by targeting the nucleoporin Ndc1 (Linder et al., 2017). There are three other transmembrane nucleoporins conserved in vertebrates, Nup210/gp210, POM121, and TMEM209, which was recently identified (Antonin et al., 2005; Kohlhause et al., 2026; Stavru et al., 2006b, 2006a). TMEM209 impacts cell cycle progression and cell proliferation and cooperates with Nup210/gp210 (Kohlhause et al., 2026). The functional importance of TMEM209 extends across eukaryotes: in *Arabidopsis*, the TMEM209 homolog PNET1 is expressed preferentially in actively dividing cells, and its cell cycle-dependent phosphorylation coordinates NPC disassembly and reassembly during mitosis (Fang et al., 2025). TMEM209 is also upregulated in hepatocellular carcinoma and proposed as a therapeutic target (Fang et al., 2024; Fujitomo et al., 2012). Despite these connections to cell division and disease, the molecular function of TMEM209 and its homologs during mitosis is unclear.

Here, we identify NPP-28, the *C. elegans* homolog of TMEM209, as a transmembrane nucleoporin that spatially regulates PLK-1 between nuclear envelope-associated and nucleoplasmic pools. *npp-28* loss reduces the nuclear envelope-associated pool of PLK-1 and increases its nucleoplasmic accumulation. In contrast, loss of other nucleoporins with identified docking sites for PLK-1 deplete PLK-1 from both compartments. Deletion of *npp-28* delays NPC and lamina disassembly, while faster nuclear accumulation PLK-1 correlates with premature anaphase onset. We identify Polo-Box Domain (PBD) binding sites in NPP-28 required to partition PLK-1 nuclear envelope and nucleoplasmic pools. We further show that NPP-28 is phosphorylated by PLK-1 *in vitro* and find that mutation of predicted phosphorylation sites results in earlier, rather than delayed, NEBD completion. Thus, NPP-28 both promotes NEBD and modulates its duration. Genetic analyses further reveal context-dependent functional relationships between NPP-28 and other transmembrane nucleoporins. Together, these findings suggest NPP-28 coordinates NEBD with anaphase onset by spatially regulating PLK-1. This mechanism may explain why TMEM209 homologues impact cell proliferation.

## Results

### B0464.6/NPP-28 is the *C. elegans* homologue of transmembrane nucleoporin TMEM209

We identified *C. elegans B0464.6* as a candidate nuclear pore transmembrane component in an exploratory high content phenotypic screen that sought to functionally classify genes associated with nuclear structure and function. In this screen of 148 genes, which was an extension of prior work (Green et al., 2011), phenotypic profiles were generated by analyzing the consequences of gene knockdown on the morphology of the *C. elegans* gonad. Genes with similar functions were identified by pairwise comparison of phenotypic profiles across the dataset. Genes whose knockdown led to similar phenotypes were identified based on their connection specificity index (CSI) score, which is a measure of the similarity and specificity of the compared profiles (described in (Green et al., 2011)) (Fig. S1A). Strong phenotypic matches were found for 13 of the 14 unnamed genes in the dataset (Table S1), which were included because they were identified as homologues of nuclear envelope-associated proteins through BLAST searches of a prior human nuclear envelope proteomics screen (Schirmer et al., 2003). Among these, *B0464.6,* the *C. elegans* homologue of TMEM209/NET31, was a close phenotypic match with *npp-22^NDC1^*, an integral membrane nucleoporin that links nuclear pore complex components to the pore membrane (Mansfeld et al., 2006; Mauro et al., 2022; Onischenko et al., 2009; Stavru et al., 2006a), but not with any of the other nuclear pore complex proteins included in the screen dataset (Fig. S1B-C).

We analyzed the sequence and structural homology of B0464.6 to TMEM209, which has recently been characterized as the fourth human transmembrane nucleoporin (Kohlhause et al., 2026). B0464.6 contains two predicted N-terminal transmembrane domains, a middle unstructured region, and a C-terminal structured domain that shares ∼50% amino acid similarity to human TMEM209. AlphaFold 3 modeling shows a similar predicted domain architecture to vertebrate TMEM209 as well as plant and trypanosomes homologs (Fig. 1A; (Abramson et al., 2024; Butterfield et al., 2024; Fang et al., 2024, 2025; Fujitomo et al., 2012; Kohlhause et al., 2026; Obarska-Kosinska et al., n.d.). Based on this conservation, B0464.6 was named *tmem-209/npp-28,* following the naming convention of nucleoporins in *C. elegans*.

**Figure 1.**
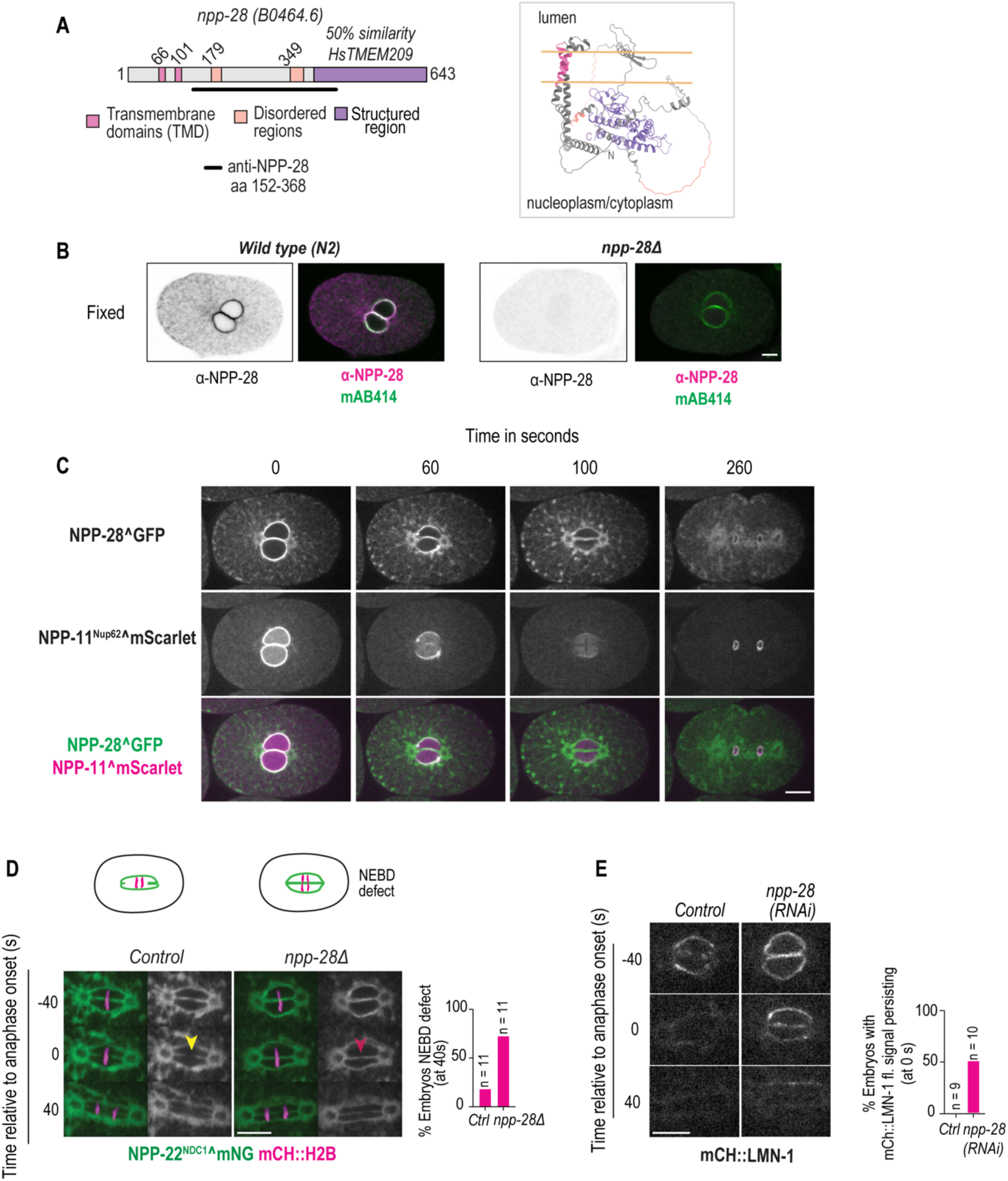
NPP-28 (C. elegans TMEM209) promotes pronuclear fusion and lamina disassembly in early *C. elegans* embryos. A. Schematic representation of NPP-28 protein domains (left). The region used to generate anti-NPP-28 antibodies is indicated by a black line. AlphaFold 3-predicted structure of NPP-28 (right). Colors according to domain organization. Peach lines indicate the predicted bilayers based on predicted transmembrane helices shown in pink. B. Fixed wild-type (N2) and *npp-28Δ* one-cell embryos coimmunostained with anti-NPP-28 and mAb414 antibodies, which in *C. elegans* recognizes NPP-10 (Mauro et al., 2022). C. Representative spinning disk confocal images of endogenously tagged NPP-28^GFP and NPP-11^Nup62^^mScarlet (Thomas et al., 2023) during the first mitotic division of *C. elegans* embryos. Time in seconds. D. Representative spinning disk confocal images of NPP-22^NDC1^^mNG and mCh::H2B embryos under the indicated conditions. Yellow arrowhead marks successful (Control) and red arrowhead marks failed (*npp-28*Δ) pronuclear membrane fusion. Plot represents percentage of embryos exhibiting pronuclear membrane fusion defects at anaphase onset (40 s). E. Representative spinning disk confocal images of mCh::LMN-1 embryos under the indicated conditions. Time is shown relative to anaphase onset in seconds. Plot shows the percentage of embryos retaining mCh::LMN-1 signal at anaphase onset (0 s). Scale bars, 10 μm. *n* indicates the number of embryos analyzed.

We characterized NPP-28 localization and dynamics during mitosis in *C. elegans* embryos. First, antibodies were raised against the central region of the NPP-28 protein (amino acids 152–368) and localization of endogenous NPP-28 was determined. NPP-28 co-localized at the nuclear rim with NPCs and to the endoplasmic reticulum (ER), as expected for a transmembrane-containing NPC-associated protein (Fig. 1B). A CRISPR/Cas9-mediated *npp-28* deletion strain was generated, which confirmed the specificity of this localization (Fig. 1B; Supplemental Fig. S1D and E). Endogenously tagged NPP-28^GFP co-localized with a member of the central channel NPP-11^Nup62^^mScarlet at the nuclear rim, which was used as a live marker for a soluble NPC component (Fig. 1C, Movie 1). While both proteins were enriched at the nuclear rim, NPP-11^Nup62^ also displayed a diffuse nucleoplasmic signal, whereas NPP-28 associated with the ER. The additional ER-associated pool is consistent with the prediction that NPP-28 is integral to the membrane, which is different from the nucleoplasmic localization of the soluble nucleoporin NPP-11^Nup62^.

We next characterized the dynamics of NPP-28 during NE breakdown (NEBD) and reformation in mitosis, using NPP-11^Nup62^ as a benchmark. Nuclear rim enrichment of NPP-28^GFP and NPP-11^Nup62^^mScarlet was progressively lost with similar kinetics as mitosis progressed (Fig. 1C). NPP-28 redistributed throughout the ER, including disassembling nuclear membranes, while NPP-11^Nup62^ appeared soluble throughout the cytoplasm and remained enriched within the mitotic spindle region (Fig. 1C). Following mitotic exit, both proteins reassociated with the reforming nuclear envelope with similar kinetics (Fig. 1C).

*npp-28Δ* embryos were viable, despite the reduced brood size, indicating that NPP-28 function at the NE is not essential for embryogenesis (Supplemental Fig. S1F), and supports germline function, in line with gonad morphology phenotypes identified in screen (Supplemental Fig. S1C and D).

Together, these data establish NPP-28 as a transmembrane ring nucleoporin, with similar nuclear envelope localization and dynamics as NPP-11^Nup62^.

### NPP-28 loss delays lamina and NPC disassembly

*npp-28Δ* embryos displayed several defects during mitosis. During mitotic prophase, mitotic chromosomes were abnormally localized to the nuclear periphery (Supplemental Fig. S1G and H), a phenotype also seen in *npp-3^Nup205^-* depleted embryos (Jiang et al., 2026). In addition, approximately half of embryos exhibited delayed NE disassembly, as assessed by pronuclear fusion failure and delayed lamina disassembly (Fig. 1D, E and Supplemental Fig. S1I). In the *C. elegans* zygote, the separate NEs surrounding the female and male pronuclei meet and undergo membrane fusion directly prior to anaphase onset to allow for merging of the chromosomal contents. Pronuclear fusion coincides with the disappearance of the nuclear lamina (Rahman et al., 2020; Velez-Aguilera et al., 2020). Pronuclear membranes marked by NPP-22^NDC1^^mNG or a general ER marker (SP12::GFP) persisted between chromosomes in a majority of *npp-28Δ* embryos as they progressed through anaphase, indicating incomplete membrane fusion (Fig. 1D and Supplemental Fig. S1I). Similarly, *npp-28* deletion delayed disassembly of nuclear lamins in many embryos, as seen by the persistence of mCH::LMN-1 signal in the chromosome and spindle region at time points in which LMN-1 was cleared in control embryos (Fig. 1E). Thus, NPP-28 contributes to NE-dependent events during mitosis.

We analyzed how loss of *npp-28* impacts the dynamics of NPC disassembly during NEBD. AlphaFold 3 predictions show NPP-28 associates with several NPC components modeled based on evidence of interactions in other systems (see Fig. 2A, B and Supplemental Fig. S2A-C; (Abramson et al., 2024; Butterfield et al., 2024; Obarska-Kosinska et al., n.d.)). These include the N-terminal transmembrane and adjacent luminal regions of NPP-28 with NPP-12^gp210^ (Fig. 2B and Supplemental Fig. S2A), its partially unstructured middle domain with the scaffold NPP-3^Nup205^ (Fig. 2B and Supplemental Fig. S2B), and its C-terminal structured domain with the central channel complex NPP-1/-4/-11^Nup54/58/62^ (Fig. 2B and Supplemental Fig. S2C). The resulting models closely resembled those predicted for human TMEM209 (Obarska-Kosinska et al., n.d.), suggesting that these interactions are conserved in *C. elegans* (Fig. 2B, Supplemental Fig. S2).

**Figure 2.**
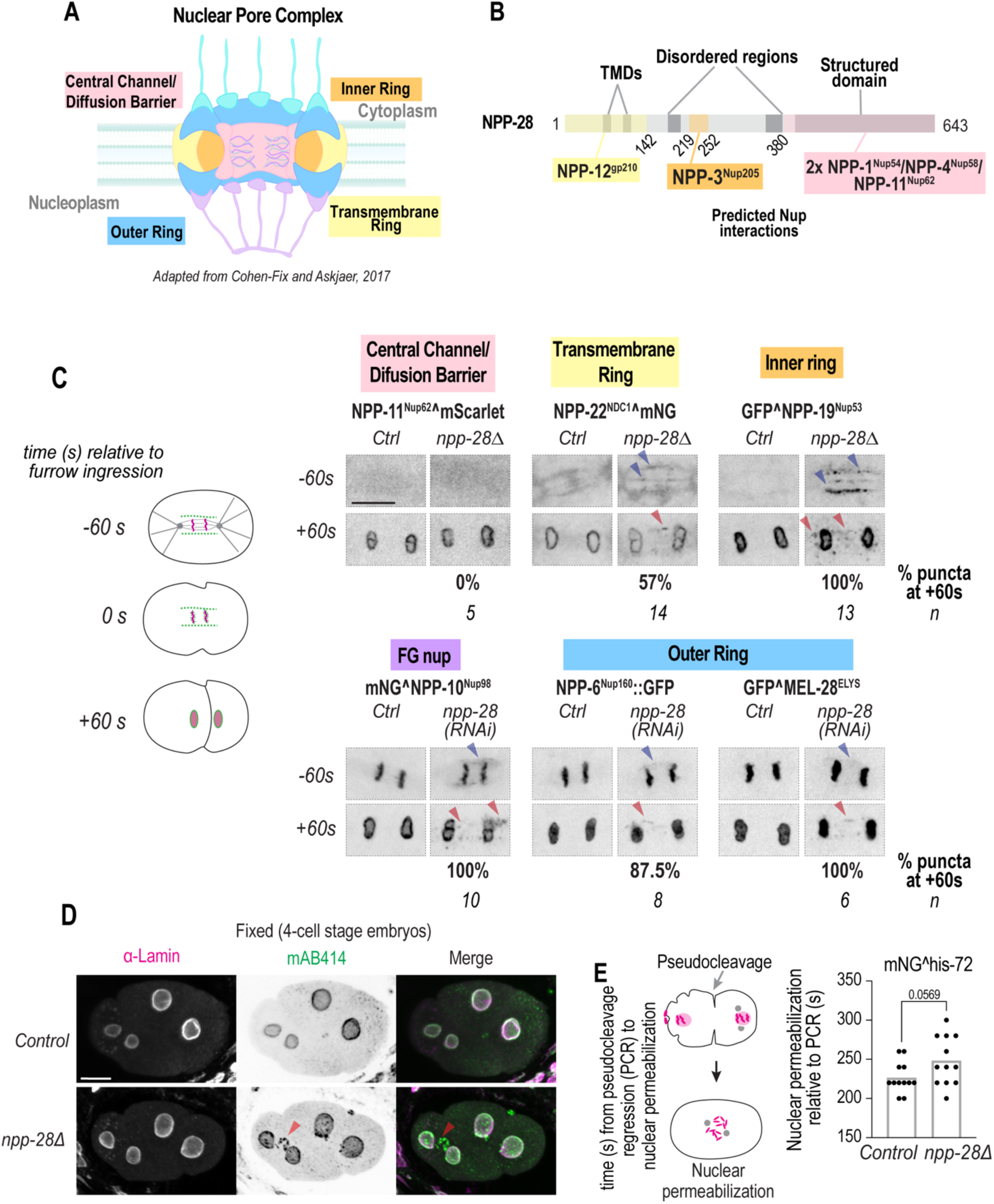
NPP-28 regulates the dynamics of multiple nucleoporin subcomplexes during mitosis. A. Schematic representation of major subcomplexes that make up different regions of the nuclear pore complex (NPC) (modified from Cohen-Fix and Askjaer, 2017). B. Schematic of NPP-28 domain organization with regions predicted to interact with indicated nuclear pore components (see Supplemental Figure 2). Colors correspond to subcomplexes highlighted in panel (A). C. Schematic of NPC dynamics relative to the onset of cytokinetic furrow ingression (0 s). Representative spinning disk confocal images of indicated fluorescently tagged nucleoporins, grouped and color-coded according to the NPC subcomplexes shown in panel A, in indicated conditions. Images of the nuclear region is shown at 60 s before and 60 s after the onset of furrow ingression (0s). Percentage of embryos exhibiting persistent nuclear envelope remnants of the indicated nucleoporin at 60 s timepoint is shown. *n* indicates the number of embryos analyzed. Blue arrowheads mark disassembly defects prior to furrow ingression and red arrowheads mark failed ectopic nucleoporin seeding sites post furrow ingression. D. Representative spinning disk confocal images of fixed embryos immunostained with mAB414 and anti-lamin antibodies in indicated conditions. To account for variable staining across slide, brightness in control was increased 3X and 4X relative to *npp-28*Δ for the mAB414 and lamin channels, respectively. Red arrowheads mark failed ectopic nucleoporin seeding sites. Scale bars, 10 μm. E. Schematic of nuclear permeabilization analysis based on loss of diffuse nucleoplasmic histone. Plot shows time in seconds of nuclear permeabilization relative to pseudocleavage regression (PCR) in mNĜHIS-72 expressing embryos in indicated conditions. Each dot represents individual embryo. p value was calculated using the Welch’s t-test with normal distributions, calculated by a Shapiro-Wilk test.

We analyzed the dynamics of representatives of NPC subcomplexes, including the outer ring scaffold (NPP-6^Nup160^::GFP and GFP^MEL-28^ELYS^), the central channel diffusion barrier (NPP-11^Nup62^^mScarlet), NPP-22^NDC1^ for the transmembrane ring, and the linker nucleoporins that participate in scaffold interactions, the inner ring complex nucleoporin GFP^NPP-19^Nup53^ and the FG-nucleoporin mNĜNPP-10^Nup98^ (Fig. 2C).

Loss of *npp-28* preferentially impaired disassembly of linker nucleoporins. NPP-10^Nup98^ and NPP-19^Nup53^ exhibited incomplete disassembly and persistent localization to nuclear envelope remnants (Fig. 2C and Movie 2). This was also confirmed in fixed embryos immunostained with mAB414 antibodies raised against FG nucleoporins (Fig. 2D), which, in worms, recognizes NPP-10^Nup98^ (Mauro et al., 2022), and was not observed for lamins, in line with our live imaging data of a delay in disassembly that led to eventual clearance between segregated chromosomes (Fig. 2D; also see Fig. 1E). Although NPC reassembly around segregated chromosomes appeared unperturbed, NPC disassembly remnants frequently seeded ectopic sites of assembly upon mitotic exit, especially at regions between segregated chromosomes and sometimes also around spindle poles (Fig. 2C, also see Movie 2). While not as dramatic, a pool of NPP-22^NDC1^, which largely redistributes to the ER upon mitotic disassembly (Mauro et al., 2022), also persisted between segregating chromosomes and, in 57% of embryos, these sites increased in signal intensity as the NE reformed upon mitotic exit, suggesting that they consist of both membranes and NPC components (Fig. 2C). Outer ring Y-complex components (NPP-6^Nup160^::GFP and GFP::MEL-28^ELYS^) did not exhibit delays in disassembly as severe as those observed for the linker nucleoporins (Fig. 2C), although fluorescent signal persisted in a majority of embryos and punctate structures appeared between chromosomes during reassembly (87.5% and 100%, respectively; Fig. 2C).

Unlike the other nucleoporins, NPP-11^Nup62^, a component of the NPC diffusion barrier, displayed normal disassembly and did not form ectopic assemblies upon mitotic exit (Fig. 2C). Nuclear permeabilization was delayed by an average of 20 s (Mean value, Control, ∼230 s; npp-28(RNAi), ∼250 s), as assessed by loss of diffuse HIS-72 nucleoplasmic signal, though this difference was not significant (Fig. 2E). The modest delay in the initiation of nuclear permeabilization in the absence of *npp-28* is consistent with the limited effect on the disassembly dynamics of NPP-11^Nup62^. Together, these findings indicate that NPP-28 preferentially promotes the timely and complete disassembly of the linker nucleoporins NPP-19^Nup53^ and NPP-10^Nup98^, with a more pronounced effect on NPP-19^Nup53^.

### NPP-28 concentrates PLK-1 at the nuclear envelope and limits its nucleoplasmic accumulation

NPP-28 concentrates Polo-like kinase 1 (PLK-1) at the nuclear envelope while restricting its nucleoplasmic accumulation. In mitotic embryos, PLK-1 localizes to spindle poles, chromosomes, and regions of the NE located at the pronuclear interface and adjacent to the spindle poles (Fig. 3A; (Chase et al., 2000; Martino et al., 2017)). Upon nuclear permeabilization, PLK-1 remains associated with nuclear membranes and accumulates within the mitotic spindle region (Fig. 3A, Movie 3). *npp-28* deletion reduced PLK-1 at the nuclear envelope throughout all mitotic phases and increased its nucleoplasmic levels, especially evident during early mitotic prophase (Fig. 3A-E, Movie 3). In contrast, *npp-1^Nup54^,* which docks and is a substrate for PLK-1(Martino et al., 2017), globally impairs nuclear transport and causes NPC clustering (Schetter et al., 2006), as also seen with NPP-28^GFP (Supplemental Fig. S3A). Consistent with this, partial *RNAi-*depletion of the central channel nucleoporin NPP-1^Nup54^ reduced PLK-1 from the NE, nucleoplasm, and mitotic chromosomes (Fig. 3A).

**Figure 3.**
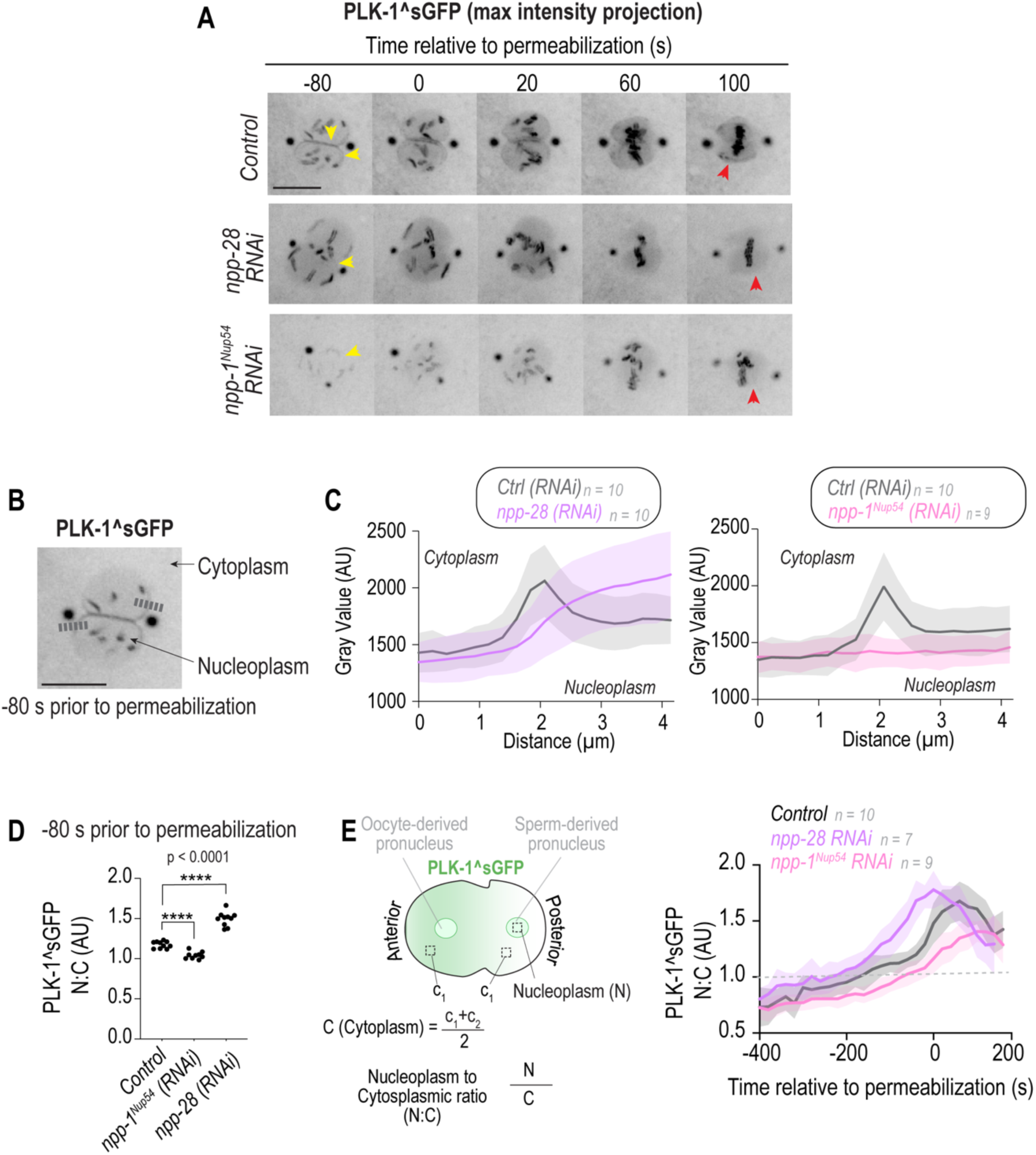
NPP-28 concentrates PLK-1 at the nuclear envelope and restricts its nuclear accumulation. A. Representative maximum-intensity projections of PLK-1^sGFP in indicated conditions. Time is shown in seconds relative to nuclear permeabilization, as determined by loss of diffuse nucleoplasmic mCh::H2B signal (not shown). Yellow arrowheads mark nuclear rim at a timepoint before nuclear permeabilization. Red arrowheads mark nuclear rim region at timepoint after nuclear permeabilization. B. Representative image of PLK-1^sGFP fluorescence 80 s prior to nuclear permeabilization showing dashed lines to indicate regions used for line-scan analysis in (C) drawn from the cytoplasm to the nucleoplasm. C. Plots of mean fluorescence intensity profiles of PLK-1^sGFP measured along line scans as shown in (B) in indicated conditions. Shaded regions represent SD. D. Plot showing quantitation of nucleoplasmic-to-cytoplasmic (N:C) ratios of PLK-1^sGFP fluorescence 80 s prior to nuclear permeabilization, as calculated from fluorescence intensity profiles in panel C, in indicated conditions. Each dot represents a single embryo. **** indicates p < 0.0001. p values were calculated using Brown-Forsythe and Welch ANOVA tests with normal distributions, as calculated by a Shapiro-Wilk test. E. Schematic illustrating quantification of the PLK1^sGFP nucleoplasmic-to-cytoplasmic (N:C) ratio over the time course prior to the first cell division. Plot shows the time course of PLK-1^sGFP N:C ratio relative to nuclear permeabilization in indicated conditions. Note that control embryos were collected from worms incubated at 20°C, while *npp-1(RNAi)* embryos were collected from worms incubated at 15°C for 24 hours prior to imaging at 20°C. Shaded regions represent SD. Scale bars, 10 μm. *n* indicates the number of embryos analyzed.

To quantify the differences in PLK-1^sGFP at the NE, we measured the mean GFP fluorescence intensity along line profiles extending from the cytoplasm to the nucleoplasm at a defined time point prior to permeabilization (-80s, Fig. 3B). In control embryos, peak fluorescence corresponded to the NE–associated pool, with a ∼1.2-fold enrichment of nucleoplasmic relative to cytoplasmic signal (Fig. 3C, D). *npp-28* RNAi-depletion abolished the discernible peak the NE-associated peak of PLK-1^sGFP and elevated its nucleoplasmic signal, resulting in a nuclear-to-cytoplasmic ratio of ∼1.5 (Fig. 3C, D). In *npp-1^Nup54^* RNAi-depleted embryos, the fluorescence intensity of PLK-1^sGFP was uniformly reduced across compartments, yielding a nuclear-to-cytoplasmic ratio of ∼1 (Fig. 3C, D).

Because NPP-1^Nup54^ is a central component of the NPC diffusion barrier and its loss causes defects in the distribution of NPCs, its effects on nucleoplasmic PLK-1 levels could reflect a general defect in nuclear permeability and NPC architecture. To test if this could also explain its effects on PLK-1 NE localization, we depleted importin-α/β (IMA-2 and IMB-1) to disrupt global nuclear import and imaged through early and later stages of mitosis. As expected, RNAi-mediated depletion of importin-α/β resulted in minimal nuclear import of PLK-1 prior to nuclear permeabilization (Supplemental Fig. S3B). Notably, under these import-deficient conditions, PLK-1 association with the nuclear rim was significantly delayed relative to controls; however, PLK-1 became enriched at the nuclear rim at time points when permeabilization normally occurs, suggesting that PLK-1 accesses docking sites only at later stages when import is defective (Supplemental Fig. S3B; (Martino et al., 2017)). Together, these data indicate that nuclear import is required for early, but not late, recruitment of PLK-1 to the NE. By contrast, depletion of either NPP-28 or NPP-1^Nup54^ impair NE-association of PLK-1 throughout all stages of mitosis.

We next examined the dynamics of PLK-1 accumulation within the nucleus throughout the first cell division. PLK-1 accumulated in the nucleus more rapidly from interphase through mitotic prophase in *npp-28* RNAi-depleted embryos (Fig. 3E, Movie 3), and there was no evidence of a global defect in nuclear import, as measured by an NLS-GFP reporter (Supplemental Fig. S3C). The amount of time required for PLK-1 to reach a nuclear-to-cytoplasmic ratio of 1 was ∼225s prior to permeabilization in *npp-28* RNAi-depleted embryos, compared to ∼175 s and ∼50 s in control and *npp-1^Nup54^* RNAi-depleted embryos, respectively (Fig. 3E). The peak enrichment of the nucleoplasmic pool of PLK-1 in *npp-28* RNAi-depleted embryos occurred at the time of nuclear permeabilization, whereas in control embryos PLK-1 continued to rise in the mitotic spindle region as embryos progressed through mitosis, reaching maximal levels ∼100 s after permeabilization. *npp-1^Nup54^*RNAi-depletion led to a significantly delayed and slowed increase in nuclear PLK-1 and at the mitotic spindle region, which peaked only after nuclear permeabilization to lower levels than control (Fig. 3E). These data indicate that NPP-28 partitions PLK-1 between the nuclear envelope and nucleoplasm, without broadly impacting nuclear import.

### NPP-28 prevents premature anaphase onset

*npp-28* loss shortened mitotic duration by ∼30 - 60s (180-200s in controls versus 140-150s in *npp-28Δ*) (Fig. 4A). In *C. elegans* embryos, local phospho-regulation of kinetochore-bound CDC-20, the activator of the Anaphase Promoting Complex/Cyclosome (APC/C), controls mitotic duration (Kim et al., 2017). PLK-1 recruits CDC-20 to kinetochores to promote its dephosphorylation, allowing CDC-20 to activate APC/C and trigger anaphase onset (Houston et al., 2023; Kim et al., 2017). Shortened mitotic duration is therefore consistent with earlier nuclear accumulation of PLK-1, which would be predicted to promote earlier CDC-20 recruitment to kinetochores. Indeed, CDC-20 accumulates earlier at chromosomes in *npp-28Δ* embryos compared to control embryos (Fig. 4B). We also determined the genetic relationship between cyclin B1 (CYB-1) and NPP-28 because CYB-1 and CDC-20 phosphorylation counteract APC/C activation to slow mitotic duration (Lara-Gonzalez et al., 2024, 2017). RNAi-depletion of *cyb-1* shortened mitotic duration by 80 - 100 s compared to control, as expected (Lara-Gonzalez et al., 2024). Mitotic duration upon combined loss of *cyb-1* and *npp-28* was similar to that of *cyb-1* depletion alone (Fig. 4C) suggesting *npp-28* and *cyb-1* function in a common pathway controlling mitotic duration. Notably, combined loss of *npp-28* and *cyb-1* delayed chromosome congression in a subset of embryos (Fig. S3D) suggesting that despite working in the same pathway for mitotic timing, they make separable contributions to chromosome congression. We conclude that NPP-28 restricts premature nuclear PLK-1 accumulation, which enables timely PLK-1-dependent recruitment of CDC-20 to kinetochores and proper timing of anaphase.

**Figure 4.**
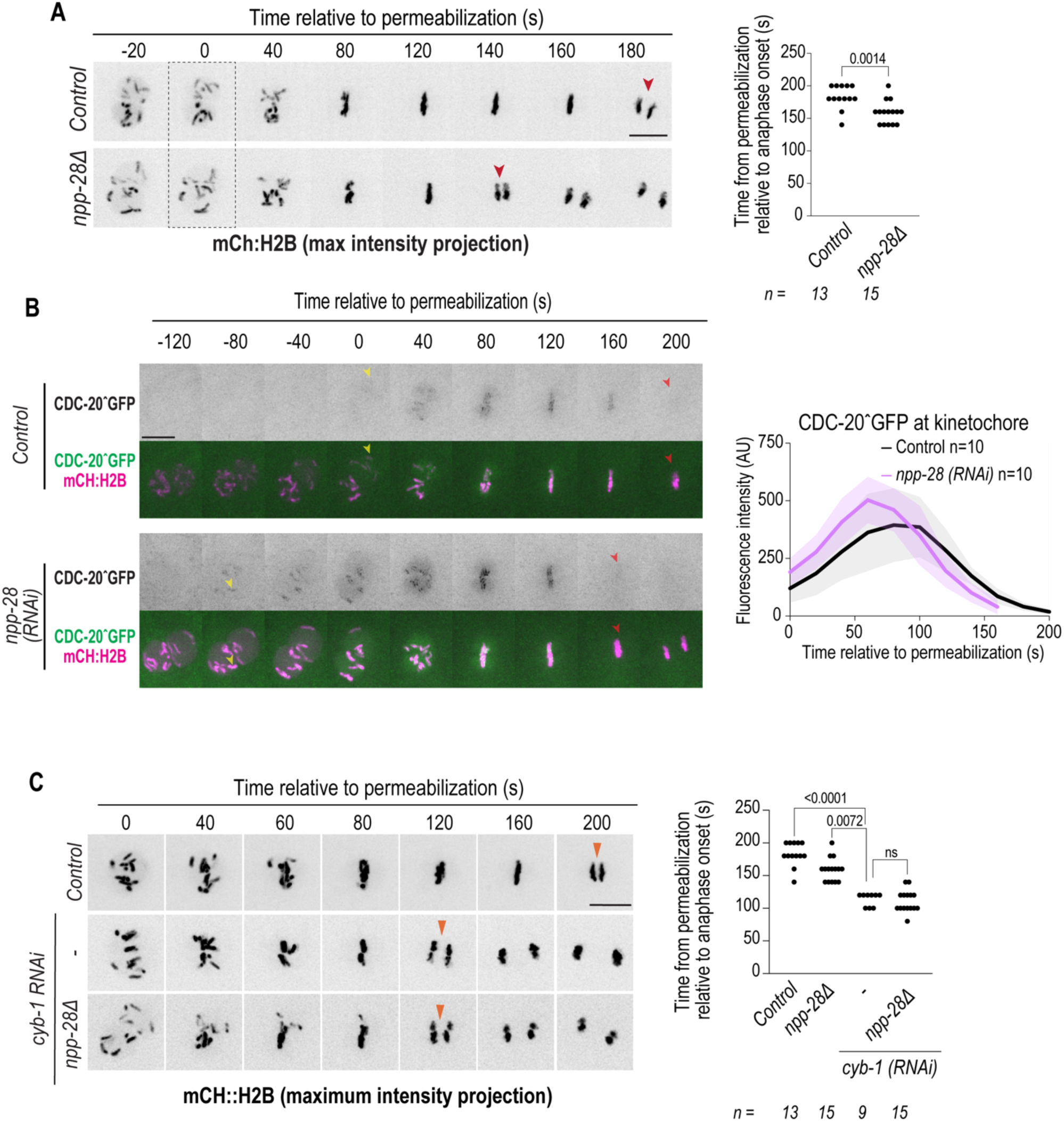
NPP-28 lengthens the duration mitosis. A. Representative maximum-intensity projections of mCh:H2B in indicated conditions. Time is shown in seconds relative to nuclear permeabilization, as determined by loss of diffuse nucleoplasmic mCh::H2B signal. Red arrowheads indicate first frame where anaphase is visible. Plot represents the time from nuclear permeabilization to anaphase onset in indicated conditions. Each dot represents individual embryo. p value was calculated using the Mann-Whitney test with non-normal distributions, calculated by a Shapiro-Wilk test. B. Representative spinning-disk confocal images of CDC-20^GFP and mCh:H2B relative to nuclear permeabilization. Yellow arrowheads indicate first frame where CDC-20 accumulation at chromosomes is visible. Red arrowheads indicate frame in which CDC-20 leaves the chromosomes. Plot shows fluorescence intensity levels of CDC-20^GFP at chromosomes over time. C. Representative maximum-intensity projections of mCh:H2B in indicated conditions relative to nuclear permeabilization. Orange arrowheads indicate anaphase. Plot represents the time from nuclear permeabilization to anaphase onset in indicated conditions. Each dot represents individual embryo. p value was calculated using the Kruskal-Wallis test with non-normal distributions, calculated by a Shapiro-Wilk test. Control and *npp-28Δ* values repeated from A. Scale bars, 10 μm. *n* indicates number of embryos analyzed from multiple independent experiments.

### Polo-box domain docking sites in NPP-28 partition PLK-1 between the nuclear envelope and nucleoplasm

NPP-28 may concentrate a pool of imported PLK-1 at the NPC, either by docking PLK-1 itself or by facilitating the delivery of PLK-1 to docking sites on other nucleoporins. We generated domain deletions in NPP-28 to determine whether it docks PLK-1. We first tested the requirement for NEBD in two truncation mutants: a middle region deletion (*ΔM*(*aa 189-359*)) and a C-terminal domain deletion (*ΔCTD*(*aa 358-643*)) (Fig. 5A). When present as the only copy of *npp-28,* WT and *ΔM* NPP-28 supported NEBD; by contrast, *ΔCTD* did not (Fig. 5B). Both GFP-tagged *ΔM* and *ΔCTD* localized to the NE when expressed as the sole source of *npp-28*, although to a slightly reduced extent compared to endogenous NPP-28^GFP (Supplemental Fig. S5A, B). Thus, the C-terminal region of NPP-28 functionally supports NEBD.

**Figure 5.**
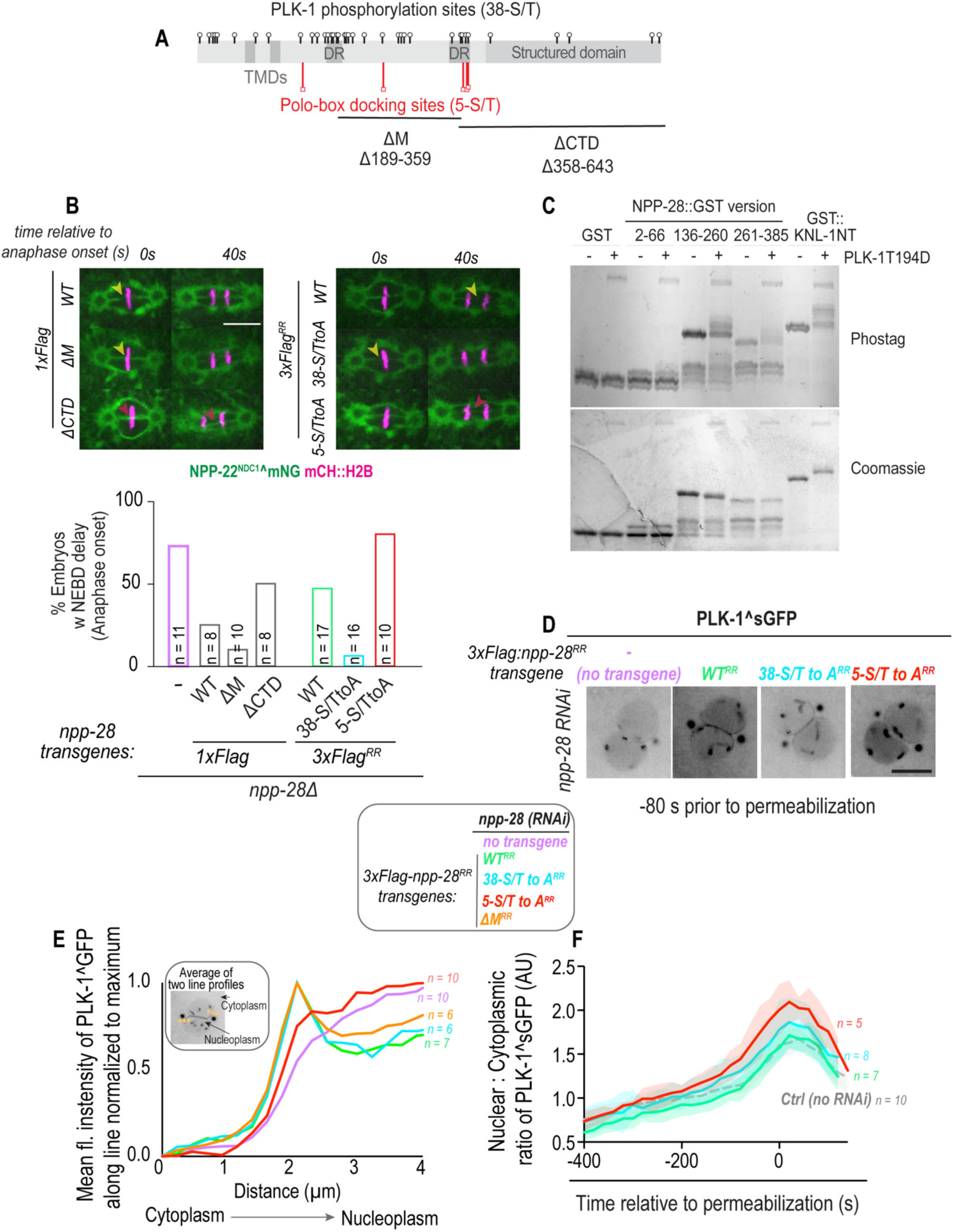
Predicted PLK-1 docking sites within NPP-28 are required for PLK-1 enrichment at the nuclear envelope, restriction of its nucleoplasmic accumulation, and timely nuclear envelope breakdown. A. Schematic of NPP-28 showing predicted PLK-1 phosphorylation sites (38-S/T), Polo-box docking (PBD) sites (5-S/T) and regions deleted in ΔM and ΔCTD constructs in relation to transmembrane domains (TMDs), disordered regions (DR), and the conserved C-terminal structured region. B. Representative spinning disk confocal images of NPP-22^NDC1^^mNG and mCh::H2B embryos under the indicated conditions. Yellow arrowheads indicate successful pronuclear fusion, while red arrowheads indicate fusion failure. Plot (bottom) of the percentage of embryos exhibiting pronuclear fusion failure at anaphase onset. C. Representative phos-tag and corresponding Coomassie stained gel of indicated GST fusion proteins incubated with or without constitutively active PLK-1, as indicated. D. Representative single slice spinning disk microscopy images of PLK-1^sGFP in indicated conditions at 80 s prior to permeabilization E. Plot of mean fluorescence intensity profiles of PLK-1^sGFP normalized from minimum to maximum values. Line profiles were measured along line scans drawn from the cytoplasm to the nucleoplasm, as shown in inset, in *npp-28* RNAi-depleted embryos expressing the indicated RNAi-resistant (RR) transgenes. F. Plot of the nucleoplasmic-to-cytoplasmic (N:C) ratio of PLK-1^sGFP fluorescence over the time course prior to the first cell division in indicated conditions. Time in seconds relative to nuclear permeabilization, as measured by loss of nuclear soluble mCh:H2B. Shaded regions represent SD. Control (no RNAi) without transgene expression is shown as dotted line duplicated from plot in Fig. 3E. *n* indicates the number of embryos analyzed. Scale bars, 10 μm.

We used Group-Based Prediction System (GPS) Polo (Liu et al., 2013; Xue et al., 2005) to identify five putative Polo Box Domain (PBD) docking sites in NPP-28 (Supplemental Fig. S4C); three of which (aa residues S368, T373, T375) overlap with the C-terminal domain truncation (Fig. 5A). Moreover, there are 38 sites that match the consensus sequence for PLK-1 phosphorylation, most of which reside in the middle, unstructured region of NPP-28 (Table S2 and Fig. 5A). Through *in vitro* kinase assays, we found that PLK-1 phosphorylated the middle regions of NPP-28, particularly the region encompassing amino acids 136-385 (Fig. 5C).

To test if PLK-1 docking sites and/or phosphorylation are important for its function in NEBD *in vivo*, we generated 3XFlag-tagged versions of NPP-28 with either 38 PLK-1 phosphorylation sites (*38-S/T-to-A*) or five PLK-1 docking sites (*5-S/T-to-A*) mutated, with only one overlapping site between constructs (aa T375, Fig. 5A and Table 1). Using antibodies against the 3XFlag tag, we detected comparable expression levels and similar localization of both mutant and WT proteins (Supplemental Fig. S4D and E). Consistent with this, the GFP-tagged PLK-1 docking mutant (*5-S/T-to-A*) also localized normally to the NE (Supplemental Fig. S4B). Interestingly, the *5-S/T-to-A* mutant recapitulated the pronuclear fusion NEBD delay observed upon *npp-28* deletion, but the phospho-deficient mutant (*38-S/T-to-A*) did not (Fig. 5B).

**Table 1:**
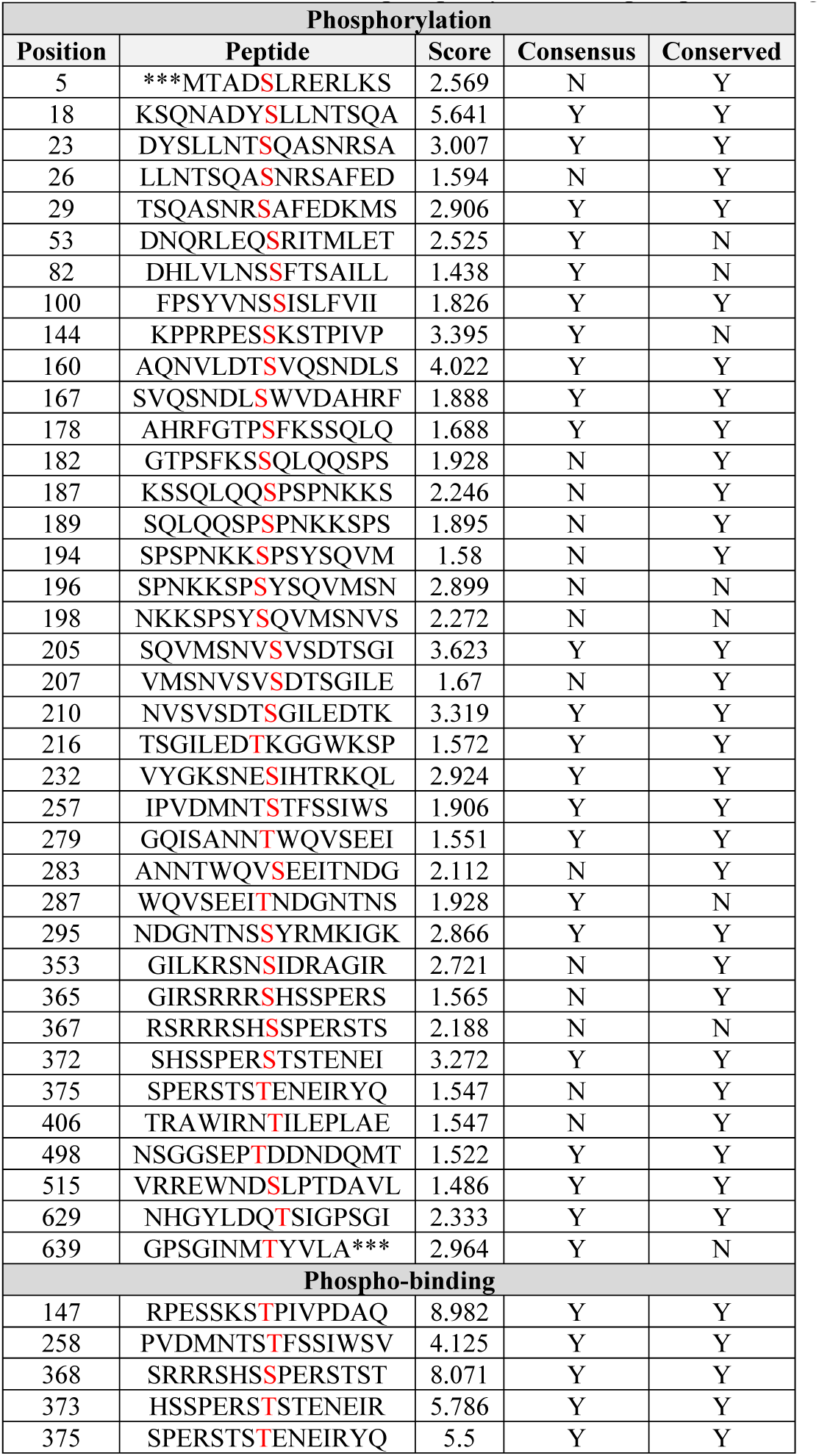
Predicted PLK-1 phosphorylation sites in NPP-28 based on Group-Based Prediction System (GPS) Polo (Xue et al., 2005) selected for mutational analysis. Amino acid position in NPP-28, peptide with consensus sequence, and rank based on GPS Polo algorithm (Xue et al., 2005) are shown. Conservation of the corresponding residues across nematode species, as determined from the multiple sequence alignment, and conformity to the established PLK-1 phosphorylation consensus motif [N/D/E/Y, X, S/T, F] are also listed, as suggested in (Houston et al., 2023). Y/N correspond to Yes/No. All sites listed in this table were chosen because they met at least two of these three criteria and, thus were selected for mutational analysis. Bottom portion of the table lists phospho-binding site candidates selected based on GPS-Polo 1.0 score and similarity to the PLK-1 polo-box domain (PBD) docking motif [F/P, F, T/Q/H/M, S, pT/pS, P/X] (Elia et al., 2003; Tavernier et al., 2015). PBD docking sites are duplicated in Fig. S4C.

Instead, the phospho-deficient mutant suppressed the pronuclear fusion defect more effectively than the WT transgene (Fig. 5B). Together, these data suggest a role for NPP-28 in directly docking PLK-1 to promote NEBD, whereas phosphorylation of NPP-28 may slow the duration of disassembly.

We next examined PLK-1 NE-docking and nuclear accumulation in the docking and phosph-deficient *npp-28* mutants. Using the same transgenic strains, nuclear rim-associated PLK-1 was quantified at 80 s prior to permeabilization and, consistent with the NEBD phenotypes, the *5-S/T-to-A* abolished PLK-1 association with the nuclear rim (Fig. 5D and 5E) whereas neither deletion of the middle region (*ΔM*) nor the phospho-deficient mutant (*38-S/T-to-A*) affected PLK-1 localization relative to the WT transgene (Fig. 5D and 5E). Measurements of nuclear PLK-1 accumulation over the time course of mitosis showed the earlier and faster nuclear accumulation upon RNAi-depletion of *npp-28* in the *5-S/T-to-A* mutant, but not the *38-S/T-to-A* or WT expressing embryos (Fig. 5F; see Fig. 3E). Both mutants suppressed the abnormal chromosome localization to the nuclear periphery resulting from loss of *npp-28* to the same extent as WT suggesting that this phenotype may be unrelated to NE accumulation of PLK-1 (Supplemental Fig. S4F). Together, these results support a model in which NPP-28 functions both as a docking platform for PLK-1 and as a substrate of PLK-1, with these roles contributing differentially to the timing of NEBD.

### NPP-28 and NPP-12^gp210^ have context-dependent and overlapping functions

NPP-28 homologs function with other transmembrane nucleoporins, particularly gp210 (NPP-12 in *C. elegans*). The two additional transmembrane nucleoporins that are conserved in *C. elegans* are NPP-22^NDC1^and NPP-12^gp210^. The *C. elegans* embryonic phenotype differs from *npp-28 -* loss of *npp-22^ndc1^*produces defects in pronuclear growth that are evident prior to NEBD complicating NEBD analysis (Mauro et al., 2022). The recovery of *npp-22 ^ndc1^* as the closest phenotypic match to *npp-28* in the gonad screen therefore suggests a potential context-dependent relationship between these transmembrane nucleoporins in *C. elegans* germline morphology. On the other hand, NPP-12^gp210^ promotes lamin disassembly during NEBD in *C. elegans* embryos (Audhya et al., 2007; Bahmanyar et al., 2014; Galy et al., 2008), suggesting it functions in the same genetic pathway as NPP-28 to promote this process.

NPP-12^gp210^ and NPP-28 were not required for the localization of the other (Fig. S5B), but did display a strong genetic interaction in both embryos and germline development. In embryos, RNAi depletion of *npp-12^gp210^* results in a high frequency of twinned nuclei at the two-cell stage following the first mitotic division, due to persistence of the nuclear lamina and failure of membrane fusion between segregating chromosomes (Audhya et al., 2007; Bahmanyar et al., 2014; Galy et al., 2008). Depletion of *npp-12 ^gp210^* in the *npp-28Δ* background resulted in a more severe NEBD defect characterized by persistent membrane connections between daughter nuclei, which remained closely apposed at time points when membrane clearance had occurred in *npp-12^gp210^*-depleted embryos alone (Fig. 6A-B, Supplemental Fig. S5A, and Movie 4). Quantification of daughter nuclei connected by membranes confirmed an exacerbation of the defect in both *npp-28Δ* embryos RNAi-depleted of *npp-12 ^gp210^* and in *npp-12 ^gp210^Δ* mutants RNAi-depleted of *npp-28* (Fig. 6B).

**Figure 6.**
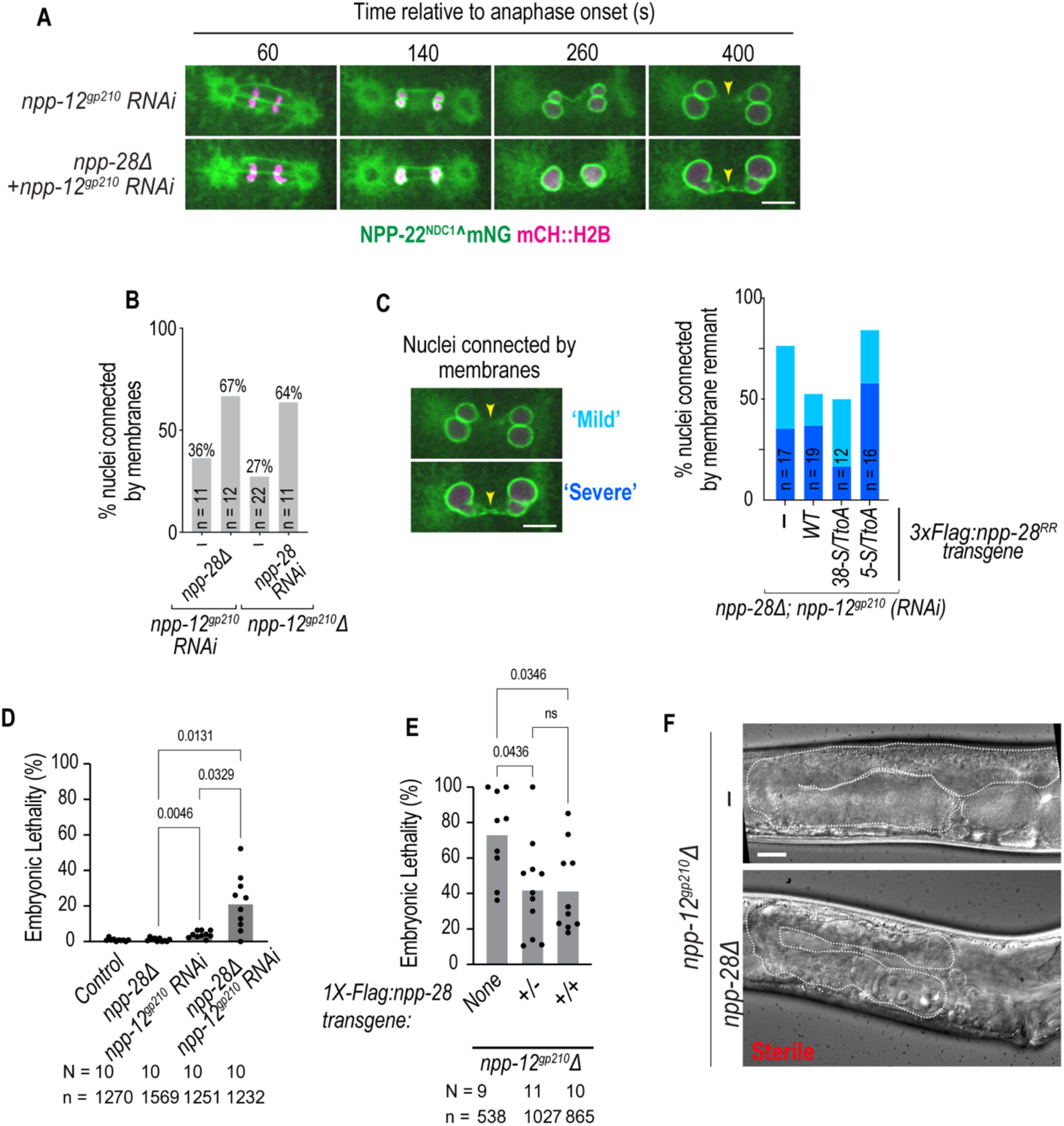
NPP-28 and NPP-12^gp210^ have partially overlapping, context-dependent roles in embryos and in germline development. A. Representative spinning-disk confocal images of NPP-22^NDC1^^mNG and mCh::H2B embryos in indicated conditions. Time is shown relative to anaphase onset in seconds. Arrowheads mark lack of membrane clearance between daughter nuclei. Scale bar, 10 μm. B. Plot of percentage of embryos containing daughter nuclei connected by persistent membrane connections at 500 s post anaphase onset following *npp-12^gp210^* RNAi depletion in *npp-28Δ* embryos or following *npp-28* RNAi depletion in *npp-12^gp210^Δ* embryos. C. Representative images of mild or severe persistent membrane connections (arrowheads) as scored by the degree of membrane signal between nuclei. Plot of quantification of mild or severe phenotypes at 400 - 500 s post anaphase onset in *npp-28Δ* embryos expressing the indicated *npp-28* transgenes following *npp-12 ^gp210^* RNAi depletion. D. Embryonic lethality of the indicated genotypes and *RNAi* conditions. Each dot represents an independent experiment. p values calculated with Brown-Forsythe and Welch ANOVA test assuming normal distributions. E. Embryonic lethality of *npp-12^gp210^Δ* worms carrying one or two copies of the WT *npp-28* transgene in the background of endogenous *npp-28*. Each dot represents an individual animal. p values calculated with Brown-Forsythe and Welch ANOVA test with normal distribution calculated by Shapiro-Wilk test. F. Representative DIC images of adult *npp-12Δ* and *npp-12Δ; npp-28Δ* worms. Dashed lines outline the germline. *npp-12 ^gp210^Δ; npp-28Δ* animals are sterile. Scale bar, 20 μm. N indicates number of worms analyzed, *n* indicates the number of embryos.

To test whether NPP-28 contributes through its role in docking PLK-1 at the nuclear envelope, we expressed NPP-28 docking and phospho-deficient transgenes in the *npp-28* deletion background, such that the transgene was the sole source of NPP-28 and performed *npp-12^gp210^* RNAi depletion. The WT NPP-28 transgene partially rescued the exacerbated NEBD phenotype, reducing the proportion of nuclei in the mild defect category, although this rescue was incomplete (Fig. 6C). However, consistent with our prediction, only the NPP-28 mutant defective in PLK-1 docking (*5-S/T-to-A*) recapitulated the severe membrane disassembly phenotype (Fig. 6C). In contrast, the phospho-deficient NPP-28 mutant suppressed the severe phenotype more effectively than the WT transgene (Fig. 6C).

Unlike RNAi-depletion of *npp-12^gp210^*, which does not result in substantial embryonic lethality (Fig. 6D), deletion of *npp-12 ^gp210^* resulted in variable embryonic lethality, with some worms failing to lay any eggs that survive to hatching, and reduced brood size (Fig. 6E and Supplemental Fig. S5C). The increased embryonic lethality observed in the deletion likely reflects roles for NPP-12^gp210^ later in embryogenesis, after zygotic transcription is activated, which are not fully captured by RNAi-mediated depletion from the germline (Oegema, 2006). Introduction of a single additional copy of *npp-28* modestly reduced embryonic lethality in *npp-12*^gp210^*Δ* worms (Fig. 6E), supporting a partial functional overlap between NPP-28 and NPP-12^gp210^ during embryogenesis (Fig. 6E). The presence of two transgenic copies of *npp-28* did not further enhance this suppression, indicating either that a single additional copy is sufficient to achieve maximal compensatory capacity or that additional copies do not substantially increase functional NPP-28 protein levels.

While only a few worms without *npp-12^gp210^* were sterile (Fig. 6E), the additional deletion of *npp-28* resulted in 100% sterility (17/17 worms) with severe defects in adult germline morphology (Fig. 6F). In almost all worms, the gonad appeared highly underdeveloped and reduced in size with no oocytes present compared to stage-matched *npp-12*^gp210^*Δ* animals (Fig. 6F). These findings reveal a strong genetic interaction between *npp-12^gp210^* and *npp-28* in the germline, extending their functional relationship beyond embryonic NEBD. The severe synthetic phenotype suggests that NPP-12 ^gp210^ and NPP-28 function in overlapping pathways whose developmental consequences are shaped by cellular context. Consistent with this idea, increasing *npp-28* copy number modestly improved embryonic viability but did not suppress the brood-size defect of *npp-12^gp210^Δ* animals (Supplemental Fig. S5C), suggesting the extent of functional overlap is context-dependent.

The 5*-S/T-to-A* transgene was significantly less effective than WT at suppressing embryonic lethality, suggesting that PLK-1 docking by NPP-28 at least partially contributes to compensation for loss of *npp-12 ^gp210^* in early embryos (Supplemental Fig. S5D). However, phospho-deficient *38-S/T-to-A* and *ΔM *NPP-28 mutants suppressed the heightened embryonic lethality resulting from *npp-12 ^gp210^ RNAi-*depletion in the *npp-28Δ* background to the same extent as the WT transgene, consistent with a lack of NPP-28 phosphorylation enhancing successful NEBD (Supplemental Fig. S5D). Notably, all NPP-28 phospho-mutants suppressed the brood-size and viability defects associated with loss of *npp-12 ^gp210^* more effectively than complete deletion of *npp-28* (Supplemental Fig. S5D and E). These results suggest that the PLK-1-dependent role of NPP-28 characterized during embryonic NEBD represents only one aspect of its functional relationship with NPP-12^gp210^, and that additional NPP-28 activities contribute to organismal development and fertility in worms.

## Discussion

NPP-28 partitions PLK-1 between the nuclear envelope and nucleoplasm to coordinate NEBD with anaphase onset. TMEM209 homologs have previously been linked to cell proliferation in both plants and human cells (Fang et al., 2024, 2025; Fujitomo et al., 2012; Kohlhause et al., 2026), but their functions in this process remained poorly understood. Whether this function is conserved and underlies the proliferation phenotypes in other systems, especially given the potential relevance of TMEM209 to cancer, is an important future direction (Fang et al., 2024; Fujitomo et al., 2012).

NPP-28 spatially regulates PLK-1 by partitioning nuclear envelope and nucleoplasmic pools during mitosis. During early mitotic prophase, NPP-28 serves as a docking platform for imported PLK-1, concentrating it at NPCs and subsequently facilitating its transfer to additional nucleoporin substrates. Among the nucleoporins examined, the disassembly of NPP-19^Nup53^ and NPP-10^Nup98^ was most severely affected by loss of NPP-28. Both proteins bind to PLK-1 through self-priming mechanisms (Nkombo Nkoula et al., 2023), consistent with a model in which NPP-28 concentrates PLK-1 to support its subsequent targeting of other nucleoporins.

NPP-28 loss increases nucleoplasmic PLK-1 without impairing global import, indicating a failure to restrict imported PLK-1 at the NE. NE-association of PLK-1 also requires NPP-1^Nup54^ in both early and late mitosis (Martino et al., 2017), but, unlike NPP-28, *npp-1^Nup54^* loss impairs global transport (Schetter et al., 2006). We suggest nuclear import initially provides PLK1 access to NPC-associated docking sites, where NPP-28 concentrates and retains PLK-1 at the NE, facilitating its transfer to additional nucleoporins, such as NPP-1^Nup54^. Later in mitosis, when the nuclear envelope is permeabilized, PLK-1 may reach NPP-28 and NPP-1^Nup54^ in an import-independent way. Our data suggest that anchoring of the imported PLK-1 to NPP-28 at the nuclear envelope early in mitosis restricts it from the nuclear interior, where PLK-1 levels must be controlled to properly regulate the timing of chromosome segregation. In support of NPP-28 regulating mitotic timing through nuclear PLK-1 accumulation, we show that *cyb-1* loss speeds up anaphase onset to a similar extent as the combined loss of *cyb-1* and *npp-28*. If *npp-28* and *cyb-1* acted independently, we would expect their combined loss to further accelerate anaphase onset. Instead, the similar timing suggests both regulate anaphase onset through CDC-20 activation, with NPP-28 acting through nuclear PLK-1 levels, while the exacerbated chromosome congression phenotypes point towards contributions from different pathways.

Our data suggest that NPP-28 promotes NEBD by recruiting PLK-1 rather than through its own phosphorylation - mutation of 38 predicted phosphorylation sites promotes NEBD to a greater extent than control contexts. The phenotypic analysis as well as results showing that NPP-28 is a substrate for PLK-1 *in vitro* supports the possibility that PLK-1 may directly phosphorylate NPP-28 to lengthen the duration of NEBD, providing a way to modulate the timing of NEBD in coordination with other mitotic events.

Our genetic analyses further highlight the importance of transmembrane nucleoporins in coordinating nuclear envelope functions with context-specific roles. The phenotypic similarity in gonad morphology between *npp-28* and *npp-22^ndc1^,* but not other genes encoding NPC components, including *npp-12* ^gp210^, provides evidence that these two transmembrane nucleoporins contribute to the same pathway in germline morphology. In contrast to the germline phenotypes, loss of *npp-22^ndc1^*in early embryos gives rise to defects in nuclear transport and reduced pronuclear size (Mauro et al., 2022), which differs from that of *npp-28*. On the other hand, NPP-12^gp210^ did not emerge as a phenotypic match in the gonad morphology screen but has a well-established role in NEBD in early *C. elegans* embryos. Furthermore, biochemical evidence in other systems suggests TMEM209 and gp210 homologs may function together. Consistent with these observations, *npp-12^gp210^* and *npp-28* genetically interact in both embryos and the germline with context-dependent functional overlap.

Context-specific functional overlap among different transmembrane nucleoporins may be a conserved feature of NPC organization across species (Antonin et al., 2005; D’Angelo et al., 2012; Galy et al., 2008; Kohlhause et al., 2026; Mansfeld et al., 2006; Mauro et al., 2022; Onischenko et al., 2009; Stavru et al., 2006a, 2006b). For example, the tissue-restricted expression and developmental functions of Nup210/gp210 in vertebrates support the idea that transmembrane nucleoporins may participate in different pathways depending on the context (D’Angelo et al., 2012; Gomez-Cavazos and Hetzer, 2015; Raices et al., 2017; Sakuma et al., 2022). How transmembrane nucleoporins cooperate or compensate for each other in different cell and tissue contexts to differentially contribute to NPC function, NE dynamics, and tissue development is an open question.

## Supporting information

Supplemental Figures and Tables with legends

## Acknowledgments

We thank members of the Bahmanyar lab for helpful discussion. We thank Jeffrey Woodruff for the gift of constitutively active PLK-1. We thank Kristin C. Gunsulus (Department of Biology, NYU) for providing input and initial computational support on pairwise comparisons for the gonad morphology screen dataset. We thank the Pintard Lab for the transgenic lamin strain. Various endogenously tagged nucleoporin strains were kindly provided by the Seydoux Lab, Askjaer Lab, and Greenstein Lab. Some strains were provided by the CGC, which is funded by NIH Office of Research Infrastructure Programs (P40 OD010440). This work was supported by grants from the NIH to S. Bahmanyar (R01 GM13004) and to P. Lara-Gonzalez (R35GM150786). Additional support was provided by the Ludwig Institute for Cancer Research to K. Oegema and American Cancer Society Postdoctoral Fellowship (PF-21-187-01-CCB) to S.R. Barger.

## Author Contributions

R.A. conducted initial experiments that were pursued by V.P., who conducted the majority of experiments.

S.R.B. generated the majority of transgenic and CRISPR-edited strains and conducted some imaging experiments and quantifications. R.A. affinity purified antibodies. T.Y.S, R.A.G, and S.B. conducted RNAi gonad morphology screen; T.Y.S generated dsRNA and collected data for the gonad morphology screen and R.A.G and S.B. analyzed phenotypes, supervised by K.O. R.B. purified the construct for antibody production.

F.M. generated a MOSCI strain and conducted some crosses. S.L. generated the PAF macro. P.L-G. advised on the design of PLK-1 mutations and conducted kinase assays. V.P., S.R.B., R.A.G, P.L-G, and S. B. worked on initial drafts and V.P., S.R.B., and S.B. finalized the manuscript, with input from all authors.

## Methods

### Strain maintenance

*C. elegans* strains used in this study are listed in Supplemental Table S1. Strains were maintained at 20°C (unless noted otherwise) on nematode growth medium (NGM) plates seeded with OP-50 *E. coli*.

### Strain generation

#### MosSCI strains

All *npp-28* constructs (re-encoded, mutant, or wild-type) were cloned into pCFJ151 for integration into the Mos I site on chromosome II (Frøkjaer-Jensen et al., 2008). Injection mixes contained the following: pCFJ151 plasmid (10 ng/μl), a plasmid encoding the Mos1 transposase (pCFJ1532, 10 ng/μl), three plasmids encoding fluorescent co-injection markers: Pmyo-2::mCherry (pCFJ90, 1.4 ng/μl), Pmyo-3::mCherry (pCFJ104, 2.9 ng/μl), and Prab-3::mCherry (pGH8, 5.7 ng/μl), and a plasmid for *peel-1* negative selection (pMA122, 10 ng/μl) (Frøkjaer-Jensen et al., 2008). Prior to injection, the mix was spun at 20,000 g for 15 min at 4°C. Adult hermaphrodites (EG8079) were injected and singled onto OP50-seeded plates until the plates were starved (∼9-10 days). Promising plates were then heat shocked for 2 hr at 34°C. The next day, non-fluorescent, wild type-moving progeny were isolated and later screened for transgene integration by PCR.

#### CRISPR-Cas9 deletion

The *npp-28* deletion strain was generated using two custom crRNAs (CRISPR guides) designed and ordered by IDT (https://www.idtdna.com/site/order/designtool/index/CRISPR_CUSTOM). crRNAs were individually incubated with trans-activating tracrRNA (IDT) for annealing (1 μl of crRNA + 1 μl of tracrRNA at 95°C for 5 min). A *dpy-10* crRNA was used for co-CRISPR selection (Paix et al., 2015). A mix of the following components was incubated at room temperature for 5 min: *[npp-28]* crRNA-1 (11.7 μM), *[npp-28]* crRNA-2 (11.7 μM), purified Cas9-NLS protein (QB3 Berkeley, 14.7 μM) and *dpy-10* crRNA (3.7 μM). Lastly, a *dpy-10* repair template (29 ng/ml) was added and the mix was spun down (20,000 g) for 30 min at 4°C. The CRISPR mix was injected into the gonads of young adult N2 worms which were singled onto individual OP50 plates. F1 progeny displaying the roller phenotype were isolated to new individual plates and, after 3 days, F1 mothers were genotyped by PCR. The deletion strain was sequenced and outcrossed 6 times to N2 worms before characterization. See Table S3 for crRNA guide sequences.

#### CRISPR-Cas9 fluorescent knock-in strains

Fluorescent endogenous tagging of *npp-28* was performed using the updated protocol by the Mello Lab (Ghanta and Mello, 2020). A gene block (Twist Bioscience) containing the fluorescent protein sequence were resuspended in nuclease-free water, melted, and subsequently cooled before adding to the mixture. The total injection mix contained 1 μl purified Cas9-NLS protein (QB3 Berkley, 14.7 μM), 3.7 μg *npp-28* crRNA-2, 100 ng/μl gene block repair template and 800 ng PRF4::rol-6(su1006) plasmid was added as a selection marker. Nuclease free water was added to a total volume of 20 μl. The mixture was centrifuged at 21,000 x g for 2 min and 17 μl was transferred to a fresh reaction tube and placed on ice until injected. Young adult N2 worms were injected and eventually singled to individual plates after 6 hours at 25°C. F1 non-rollers were genotyped for heterozygous edited mutants and singled for homozygotes. Homozygotes were sequenced and outcrossed four times to N2 worms before use.

### RNAi-based phenotypic screen

The phenotypic screen was performed as previously described (Green et al., 2011). Briefly, dsRNA was produced for 148 target genes (NS1–168; Note that numbering is non-consecutive due to omitted genes; see Table S1). Templates for dsRNA production were generated by PCR amplification of 500–1000 bp regions from genomic DNA using primers bearing T3 and T7 promoter tails (Table S1). Primer pairs amplified regions within 500–1000 bp of the target range. NS1–96 primers were ordered as a 96-well plate (Bioneer); NS125– 153 were designated interface genes for which dsRNA from the original Green et al. screen was used directly. These genes are from functionally diverse categories — including spindle assembly checkpoint, cytoskeletal, and splicing factors — selected to provide phenotypic reference points for interpreting NPC/NE gene profiles and broaden the phenotypic landscape of the dataset. For NS112–120 Sönnichsen et al. (Sönnichsen et al., 2005) primer sequences were retrieved from PhenoBank (https://phenobank.mpi-cbg.de/). NS121-124 and NS157–161 primer sequences are from (Moyle et al 2014, Gerson-Gurwitz, 2016). PCR products were column-purified (PerfectPrep 96 kit, Eppendorf) and used as templates for T3 and T7 in vitro transcription (MEGAscript T3 and T7 kits, Ambion). Sense and antisense transcription reactions were mixed, purified (MegaClear96 kit, Ambion), and annealed in 1× soaking buffer (32.7 mM Na₂HPO₄, 16.5 mM KH₂PO₄, 6.3 mM NaCl, 14.1 mM NH₄Cl) at 68°C for 10 min followed by 37°C for 30 min.

For RNAi delivery, L4-stage worms from strain OD95 were rinsed in M9 and soaked in 5 μl dsRNA supplemented with 0.5 μl of a 1:1 mixture of 63 mM spermidine and 1.1% gelatin for 24 hr at 20°C in a humid chamber. Worms were then transferred to NGM plates seeded with OP50 E. coli and allowed to recover for 48 hr at 20°C prior to imaging.

Three regions of the gonad (distal tip, pachytene and turn regions) were imaged in strain OD95 by spinning disk confocal microscopy on a Andor Revolution XD Confocal System (Andor Technology) with a confocal scanner unit (CSU-10; Yokogawa), mounted on a TE2000-E (Nikon) equipped with a 60×/1.4 Plan-Apochromat objective, solid-state 100 mW lasers, and an electron multiplication back-thinned charge-coupled device camera (iXon; Andor Technology). For each worm, a z-series of 80 planes (0.5 μm steps) was collected in the turn region and pachytene regions and 40 planes (1 μm steps) for distal tip region, encompassing DIC, GFP, and RFP channels. Phenotypic profiles were scored for each gene using a binary descriptor set (Table S1) for all three imaged regions and compared pairwise across all genes using the connection specificity index (CSI), which quantifies the similarity and specificity of phenotypic profiles between gene pairs (Green et al., 2011). Complete phenotypic scoring was obtained for 148 genes (Table S1). Pairwise CSI values were calculated and rank ordered for each gene to determine matching genes with the closest phenotypic profiles; these are listed in Table S1 as “top matches”. “Reciprocal matches” represent the highest-confidence phenotypic matches where the query gene and the ’hit’ are in each other’s top 10 phenotypic matches, based on rank-ordered CSI (Table S1). Reciprocal matched genes are identified preliminarily as candidates in the same complex or pathway and are predicted to have similar functions.

### RNAi – *C. elegans* embryo experiments

Primers for dsRNA production for *C. elegans* embryo experiments are listed in Table S2. Primer sequences were obtained from PhenoBank (https://phenobank.mpi-cbg.de/) or designed using Primer3 (https://primer3.ut.ee/) to amplify a specific region of the gene of interest from N2 gDNA or cDNA template. A T7 enzyme (MEGAscript, Life Technologies) was used to reverse transcribe the amplicon following column purification. Phenol-chloroform and isopropanol were used to purify the resulting RNAs which were then re-suspended in soaking buffer (32.7 mM Na_2_HPO_4_, 16.5 mM KH_2_PO_4_, 6.3 mM NaCl, 14.2 mM NH4Cl). After re-suspension, single stranded RNAs were annealed at 68°C for 10 min, followed by 37°C for 30 min. Double-stranded RNAs (dsRNAs) were diluted to a final concentration of ∼2000 ng/μl and stored as 2μl aliquots at −80°C. A fresh aliquot diluted to ∼1000 ng/μl with soaking buffer and centrifuged at 20,000 g for 30 min at 4°C was used for each experiment. 0.35 μl of the diluted dsRNA was loaded into the back of a pulled capillary needle and injected into the gut of L4 worms. After injection, worms were allowed to recover on OP50 seeded plates prior to imaging or brood size/lethality assays.

Knockdown time for the following targets was as follows: *npp-28*– 24 hr at 20°C, *npp-1^Nup54^* – 24 hr at 15°C, *npp-12^gp210^* – 24 hr at 20°C, *ima-2*, *ima-3* and *imb-1*– 18h at 20°C, *cyb-1*– 24 hr at 20°C, *npp-28 (*used to deplete endogenous *npp-28* in transgenic strains) – 48h at 20°C.

### Lethality and brood size quantification

L4 worms were singled 24 hours at 20°C. Worms were then singled to individual plates for 24 hours (day 1). Worms were transferred to a second plate for a final 24 hours (day 2). Worms were then killed and plates corresponding to 24-48 hours post-injection (day 1) were counted for hatched larvae and unhatched embryos. The next day the plates corresponding to 48-72 hours post-injection (day 2) were counted. The total number of embryos and larvae were combined for each time window to calculate the brood size, and the 48-72 hours embryonic lethality is reported. Conditions that did not require injections were assayed for an extra 24h, as they do not require initial recovery. Sterility of *npp-28*; *npp-12^gp210^* double mutants was confirmed across three generations, with more than 50 animals singled and scored in total.

### Generation of NPP-28 antibody

A GST-tagged NPP-28 was cloned and expressed in BL21(DE3) Competent *E. coli*. Blood was collected from rabbits injected with purified GST-NPP-28, processed to obtain serum and stored at -80 C°. Purified GST and GST-NPP-28 were desalted into coupling buffer (50 mM HEPES pH 8.0, 150 mM NaCl, 80 mM CaCl_2_, 10% Glycerol) using the BioRad Econo-Pac 10DG Disposable Chromatography Columns. Each purified protein was coupled to a 1 mL GE HiTrap NHS-Activated HP affinity column using the manufacturer-provided peristaltic pump recirculation protocol. The serum (PBS pH 7.4, 0.1% NaN_3_) was recirculated O/N through the GST-coupled column, followed by O/N recirculation over the GST-NPP-28-coupled column. The column was washed with Wash Buffer (PBS pH 7.4, 500mM NaCl, 0.1% Triton X-100) for 2 hours, followed by PBS pH 7.4 for 30 minutes. The column was washed with 10mM Tris pH 8.0 for 15 minutes. The antibody was eluted by washing with 0.1M glycine pH 2.6 and collecting fractions. The column was washed with 10mM Tris pH 8.0 for 15 minutes and additional fractions were collected after elution with 0.1M Triethylamine pH 11.5. Antibody fractions were neutralized immediately with 2M Tris pH 8.5 and the OD_280_ nm was determined. Fractions were pooled together and dialyzed against PBS pH 7.4, followed by PBS pH 7.4, glycerol 55%. The concentration was determined by measuring OD_280_ nm and aliquots were stored at -80 C°.

### Immunoblots

#### Generation of whole worm lysate

For each condition, 100 adult worms were collected in an Eppendorf tube containing 1 mL of M9 + 0.1% Triton-X and allowed to incubate for 1 hour to facilitate expulsion of ingested bacteria. Worms were then washed three times with 1 mL of M9 + 0.1% Triton X-100 (200 x g, 2 min). After the final wash, samples were brought up to a final volume of 20 μl using M9 + 0.1% Triton. Then, 20 μl of 2X Laemmli sample buffer was added, and the tubes were gently mixed. The samples were then sonicated at 70°C for 10 min, followed by heating at 95°C for 5 min. Samples were re-sonicated at 70°C for an additional 10 min. Worm lysates were stored at -20°C until gel electrophoresis.

#### Gel electrophoresis and protein transfer of worm lysates

The gel was run at 120V for 15 min to fully collapse samples and then 180V. Transfer to Nitrocellulose (BioRAD) was performed at 4°C at 900 mA for 16 hours. Membranes were blocked in Every Blot Blocking Buffer (BioRAD) for one 10 min at room temperature and incubated for 2 hours at room temperature or overnight at 4°C with the following primary antibodies diluted in blocking reagent: 1 μg/mL mouse α-alpha-tubulin (DM1A, EMD Millipore), 1 μg/mL rabbit α-B0464.6 and 1 μg/mL mouse α-Flag (Anti-Flag M2, Sigma-Aldrich), respectively. Next, membranes were briefly rinsed in TBS followed by four five-minute washes in TBST. Membranes were then incubated with the appropriate secondary antibodies for 1 hour at room temperature. Secondary antibodies were diluted 1:10,000 for horseradish peroxidase (HRP)-conjugated goat-anti-rabbit and HRP-conjugated goat-anti-mouse (Thermo Fischer Scientific). Membranes were again briefly rinsed in TBS followed by three 5 min washes in TBST. Membranes were incubated with Clarity Max Western ECL Substrate (Bio-Rad 1705062S) and ECL Prime Western Blotting System (Cytiva RPN2232), respectively, for 5 min before imaging (BioRad ChemiDoc MP Imaging Systems).

### Immunofluorescence

#### C. elegans embryos

Glass slides (Fisherbrand Premium Superfrost Microscope Slides, Fisher Scientific 12-544-7) were coated with 50-100 μg/mL poly-L-lysine (Fisher Scientific ICN15017710) and dried on a heat-block. Slides were then baked at 95°C for 30 min and then cooled to room temperature. ∼20 adult worms were picked into a 4 μl drop of ddH_2_O and gently crushed using a coverslip to express embryos from the adult worms. Slides were then immediately placed in liquid nitrogen to crack the eggshell and permeabilize the embryos. Once removed from liquid nitrogen, coverslips were quickly removed using razor blade. Slides were then fixed in cooled methanol at -20°C for 20 min. Following fixation, slides were washed two times in 1X PBS at room temperature for 10 min each using a coplin jar. After the second wash, samples were blocked with 1 % BSA/PBS per slide in a humid chamber for 1 hour at room temperature. Slides were then incubated overnight at 4°C with primary antibodies diluted in PBS (45 μl per slide; rabbit α-B0464 (1 μg/mL), mouse α-MAB414 (1 μg/mL; Biolegend). The next day, slides were washed three times in 1X PBS+ 0.2% Tween 20 (PBST) at room temperature for 10 min each using a coplin jar. Following the second wash, slides were incubated at room temperature for one hour in the dark with secondary antibodies, Goat anti-rabbit-AlexaFluorPlus-488 (Thermo scientific A32731), goat anti-mouse-AlexaFluorPlus-555 (Thermo scientific A32727) and 1 µg/mL Hoechst diluted in PBS. Slides were again washed three times in PBST at room temperature for 10 min each in the dark using a coplin jar. Mounting media (Molecular Probes ProLong Diamond Antifade Reagent) was added to each sample and coverslips were added on top and gently sealed with clear nail-polish. Slides were dried at room temperature overnight and stored at –20°C.

### Microscopy

#### Live-cell imaging of C. elegans embryos

2% agarose imaging pads were made by sandwiching molten agarose (95°C) on a glass slide. Gravid adult hermaphrodites were dissected in M9 using G10 beveled needles on a glass slide. Select embryos were transferred to the imaging pad using a mouth pipette and delicately positioned using an eyelash tool. A glass coverslip was placed to the top of the imaging pad. Imaging was performed on an inverted Nikon Ti inverted microscope with a 60X (1.4 NA) Plan Apo objective lens, a Yokogawa XI confocal scanner unit with solid state 150-mW 488 nm and 100-mW 560 nm lasers, and a CCD camera (Hamamatsu ORCA-3CCD). Typically, images were acquired every 20 seconds at five 2 μm-step z-slices in a temperature-controlled room (20°C).

To increase signal to noise ratio for line profile analysis, PLK-1^sGFP imaging of embryos expressing NPP-28 re-encoded transgenes was taken at 500ms exposure time (Fig 5D and E). For all other PLK-1^sGFP imaging (including Fig. 5F), images were taken at 200ms exposure.

#### Fixed imaging of immunostained C. elegans embryos

Fixed sample imaging was conducted on an inverted Nikon Ti2 Eclipse microscope equipped with solid state 405, 488, 561, and 640 nm lasers, a Yokogawa CSU-W1 confocal scanner unit, a 60X (1.4 NA) Plan Apo Lambda objective lens, and an sCMOS camera (Photometrics Prime BSI). 3D images were acquired with a step size of 0.3 μm.

### Image analysis

#### Peripheral Association Factor

For measuring Peripheral Association Factor or PAF of mitotic chromosomes, paternal pronuclei expressing mCh:H2B in 1-cell stage embryos were cropped at the central plane from -500 s until permeabilization.

Using Fiji, a custom-built macro was designed to measure chromatin distribution within the two regions of the nucleus. First, a whole-nucleus ROI was generated using a Gaussian blur filter (s=2) and thresholded using the Li method. Next, the original H2B image was background-subtracted, and the mean fluorescence intensity was calculated using the whole-nucleus ROI. Then, using the centroid of the whole-nuclei ROI, a smaller circle, 1/3 the area of the whole-nucleus ROI, was drawn inside to represent the nuclear center and the mean fluorescence intensity was calculated. The chromatin signal in the nuclear periphery was calculated by subtracting masked measurements of both regions. The final chromosome localization ratio is calculated by dividing the (peripheral chromatin signal multiplied by its area) by the (nuclear center chromatin signal multiplied by its area). All measurements and masking were screened for accuracy and excluded for poor segmentation. All PAF imaging and measurements were done on strains expressing mCH::H2B and NPP-22^NDC1^^mNG as markers. Additionally, PAF analysis of embryos containing transgenic NPP-28 versions were done in an *npp-28* Δ background.

#### Quantification of pronuclei fusion failure, lamina disassembly delay, chromosome congression failure, and membrane remnants

The fusion between the maternal and paternal pronuclei membranes was considered to have failed if the gap in the membrane interface between the two interfaces had not formed at the time of anaphase onset and the lingering membrane persisted past 40 s after anaphase onset. Similarly, delay in lamina disassembly was considered present if mCH::LMN-1 signal was still distinguishable at the time of anaphase onset. Chromosome congression defects were scored in the frame prior to anaphase onset. To capture all chromosomes, the mCherry::H2B channel was displayed as a maximum intensity projection of the full z-stack. Embryos were assigned to one of three categories. In “standard” embryos, all chromosomes were aligned in a compact metaphase plate oriented perpendicular to the anterior-posterior axis. In “mild” embryos, chromosomes formed a recognizable metaphase plate, but one or a few chromosomes deviated from the linear alignment. In “severe” embryos, the majority of chromosomes failed to congress to the metaphase plate by anaphase onset. Persistent membranes connecting daughter nuclei in mitosis were assessed across all z-slices at 500 s post-anaphase onset. Severity (mild/severe) of the phenotype was judged by the amount of membrane or NPP-22^NDC1^^mNG signal present between the nuclei, as well as the resulting tilt of the daughter nuclei. A mild phenotype exhibited little to no membrane between daughter nuclei, and the nuclei were not tilted toward each other.

#### PLK-1 Localization

For line scans, two 3-pixel-wide, 4-µm-long lines were drawn across the nuclear envelope of the maternal or paternal pronuclei at -80s from permeabilization. The average fluorescence intensity value among those lines was plotted after subtraction of background, measured as the average intensity of a line of the same dimensions drawn in a background region. The mCh::H2B channel was used as a reference that the center of the line was indeed at the nuclear envelope.

Normalization was performed by averaging line scans across all embryos to generate a single mean profile. The mean profile was then normalized by first dividing by its maximum value, then subtracting the minimum value, and finally dividing again by the maximum of the resulting profile, yielding a normalized line scan ranging from 0 to 1.

The nuclear-to-cytoplasmic ratio at −80 seconds prior to permeabilization was calculated by dividing the mean intensity of the five most distal values (nuclear portion) by the mean intensity of the five most proximal values (cytoplasmic portion) of the 4-μm-long line scans prior to normalization.

To measure nuclear-to-cytoplasmic ratio of PLK-1^sGFP over time, the average nuclear and cytoplasmic signal intensities were measured by drawing 5x5 pixel ROIs at each respective location for every time point. Cytoplasmic signal specifically was calculated as an average of 2 ROIs (left and right) due to gradient distribution of PLK-1. The average camera background was then subtracted from these values for each time point. The same method was used for GFP:NLS-LacI nuclear-to-cytoplasmic ratio analysis, with only one cytoplasm measurement being taken.

#### CDC-20 Localization

Chromosome-associated CDC-20s signal, indicating kinetochore localization, was quantified using a custom Fiji/ImageJ macro applied to time-lapse images acquired every 20s. Measurements were taken at every time point, from nuclear envelope permeabilization to anaphase onset. For each frame, a region of interest was selected, and chromatin was segmented in the mCH::H2B channel to mark kinetochores. The image was smoothed with Gaussian Blur (s=0.4) and thresholded using the Huang method. The resulting mask was dilated twice to include kinetochore localization, and touching chromosomes were separated by watershed segmentation. The binary mask was then multiplied by the CDC-20^GFP channel, and integrated density was measured both outside the mask and across the region of interest. Background intensity was calculated as the mean CDC-20 signal in the region outside the chromatin mask. This value was multiplied by the mask area and subtracted from the integrated density within the mask. The background-corrected integrated density was divided by the mask area to give the mean chromosome-associated CDC-20 intensity at each time point.

### Protein Sequence Analysis

Protein sequences were retrieved from WormBase and UniProtKB. Multiple sequence alignments were performed using Clustal Omega (Madeira et al., 2024) employing default parameters. Sequence similarity was visualized, and figures were prepared using ESPript 3.0 (Robert and Gouet, 2014).

Putative transmembrane helices and intrinsically disordered regions of NPP-28 were predicted using DeepTMHMM - 1.0 (Hallgren et al., 2022) and MobiDB (Piovesan et al., 2025), respectively.

Percent identity between TMEM209 and NPP-28 was calculated as the number of identical residues divided by the total number of residues. Percent similarity included conserved as well as identical residues. Two domain deletion constructs were generated: one lacking the middle region (ΔM) containing most of the disordered regions of NPP-28 (aa 189–359), and one lacking the C-terminal (ΔCTD) region (aa 358–643).

#### AlphaFold 3 complex modeling

Structural models were predicted using AlphaFold 3’s online server (https://alphafoldserver.com; (Abramson et al., 2024)). Full-length and specific residue ranges of NPP-28 (aa 1–142, 200–300), NPP-12 (aa 1723– 1796), NPP-8, NPP-3, NPP-1, NPP-4, and NPP-11 were used in the modeling. For NPP-28 (aa 1–142) and NPP-12 (aa 1723–1796) modeled with NPP-8, 12 molecules of oleic acid were included as ligand inputs to probe the predicted positioning of the NPP-28 and NPP-12 transmembrane domains relative to a lipid interface and to each other. Each predicted complex output 5 models. The highest-ranked model, based on interface predicted template modeling (ipTM) and predicted template modeling (pTM) scores, was selected for further analysis. Model confidence was assessed using predicted aligned error (PAE) plots generated by Alphafold 3. Structural models were visualized and analyzed using ChimeraX (Pettersen et al., 2021).

### PLK-1 Phospho-binding and Phosphorylation sites

Predicted PLK-1 phosphorylation and phospho-binding sites on NPP-28 were identified using GPS-Polo 1.0 (Houston et al., 2023; Liu et al., 2013; Xue et al., 2005) at the lowest threshold setting. Candidate sites were further evaluated against three criteria: a GPS-Polo 1.0 score above 1.5, conservation of the corresponding residues across nematode species as determined from the multiple sequence alignment described above, and conformity to the established PLK-1 phosphorylation consensus motif [N/D/E/Y, X, S/T, F]. Sites meeting at least two of these three criteria were selected for mutational analysis. Phospho-binding site candidates were selected based on their GPS-Polo 1.0 score and similarity to the PLK-1 polo-box domain (PBD) docking motif [F/P, F, T/Q/H/M, S, pT/pS, P/X] (Elia et al., 2003; Tavernier et al., 2015). The full list of sites is available in Table 1.

Two mutant constructs were designed in-house: one in which all 38 predicted S/T phosphorylation sites were substituted with alanine, and one in which the 5 selected phospho-binding sites were substituted with alanine. The constructs were synthesized by Twist Bioscience and subsequently cloned into appropriate vectors for the generation of transgenic MosSCI strains expressing a phospho-null NPP-28 and a docking mutant NPP-28, respectively.

### *In vitro* kinase assays

#### Protein expression and purification

cDNA fragments encoding for NPP-28 amino acids 2-66, 136-260 and 261-385 were cloned into a pET21a-based vector for their expression in bacteria as N-terminal fusions to GST. Constructs were transformed into *E. coli* BL21(DE3)pLysS cells and proteins were expressed using 0.4 mM Isopropyl β-D-1-thiogalactopyranoside (IPTG; Research Products International). Cells were collected by centrifugation and lysed in a buffer containing 25 mM Hepes pH 7.5, 300 mM NaCl, 1 mM MgCl^2^, 0.1% Triton X-100, 10 mM βME and 1 mg/mL Lysozyme, for 10 min on ice, followed by sonication. Lysates were cleared by centrifugation incubated with Glutathione Agarose resin (Pierce), for 2 hour at 4°C. GST fusion proteins were eluted from the beads with buffer containing 50 mM Tris-HCl pH 8.0, 150 mM NaCl and 10 mM reduced glutathione (TCI America). Fractions containing the highest amount of protein were pooled, dialyzed against storage buffer (25 mM Hepes pH 7.5, 200 mM NaCl, 1 mM MgCl^2^, 0.1% Triton X-100, 10 mM βME, 5% glycerol) and stored at -80°C.

#### Kinase assays

3 µg of purified protein was bound to 10 µL of Glutathione Agarose beads, for 2 hour at 4°C. Beads were washed twice with kinase buffer (25 mM Hepes pH 7.5, 50 mM NaCl, 10 mM MgCl^2^, 0.2 mM ATP, 0.1% Triton X-100, 1 mM βME) and incubated with 100 nM of constitutively active PLK-1 (T194D, a gift from Jeff Woodruff) for 1 hour at room temperature in an end-over-end rotator. Reactions were stopped by the addition of 2X sample buffer. Samples were subsequently loaded onto 10% SDS-polyacrylamide gels containing 100 µM MnCl2 and 50 µM Phos-tag (Fujiflm Wako) and stained with Coomassie blue.

Phosphorylation was assessed by the appearance of slower migrating bands.

## Statistical analysis

All data points are reported in graphs and data analysis was performed using GraphPad Prism 10. Statistical analysis was performed on datasets with multiple samples and independent biological repeats. A Brown-Forsythe and Welch ANOVA test was used to compare datasets without a normal distribution, determined by a Shapiro-Wilk test. A non-parametric Welch’s t-test was applied when comparing two datasets. A Kruskal-Wallis test was used with non-normal distributions. The type of test used, sample sizes, and P values are reported in figure legends and supplemental figure legends (p < 0.05 defined as significant).

