## Supplemental Figures and Tables with legends for "TMEM209/NPP-28 partitions PLK-1 between the nuclear envelope and nuclear interior during mitosis"

**Supplemental Figures index**

Supplemental Figures and Figure legends S1-S5

Supplemental Movie Legends Movie 1-4

Supplemental Table S1 Legend

Supplemental Tables S2-S4

### Supplemental Figures and Legends

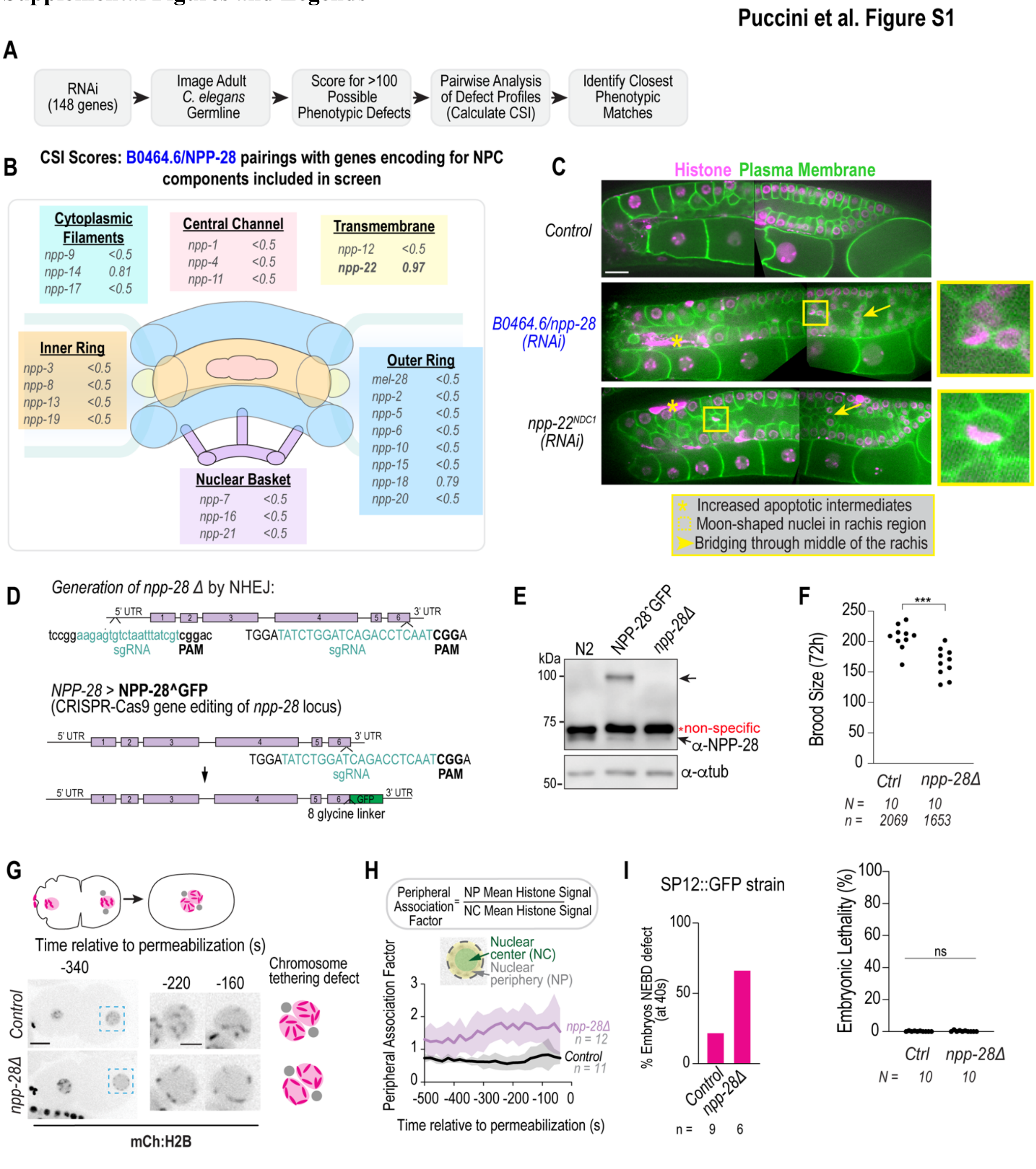


**Figure S1. Identification and characterization of B0464.6 as a putative transmembrane nuclear pore complex protein.** A. Scheme depicts the general workflow of the high content screen. Briefly, 148 genes were selected for inclusion in the screen with the goal to functionally classify all of the genes that might be associated with nuclear structure and function. For this screening effort (see Table S1), candidate nuclear envelope associated genes were identified for inclusion (Schirmer et al., 2003), along with genes associated with lamin (10), nuclear pore (23), nuclear import/export (8), nuclear envelope (6), peripheral ER (4), meiosis (9), mitosis (12), RNAi machinery (4), and an additional 28 genes that were included to help interpret and connect phenotypic scoring data to a previous screening effort (Green et al., 2011). Profiles were compared pairwise among all of the other target genes in the dataset, using the connection specificity index (CSI) metric, and the genes with the closest phenotypic match (values closest to 1) were identified. B. Nuclear pore schematic includes the *C. elegans* genes that were profiled in this work, classified according to known pore subdomain; each gene is listed with the CSI score that was calculated by comparing the profile for the uncharacterized gene, *B0464.6*$^{npp\text{-}28}$*,* with each pore-specific gene. The strongest phenotypic match for *B0464.6* was observed with the transmembrane component *npp-22*$^{ndc1}$ (CSI=0.97); weaker matches were observed with *npp-18*$^{Seh1}$ (Y-complex, CSI=0.79) and *npp-14*$^{Nup214}$ (cytoplasmic filament component, CSI=0.81), and none of the other nuclear pore components showed any appreciable phenotypic match (CSI <0.5). C. Representative images show example germlines for control, *B0464.6*$^{npp\text{-}28}$*(RNAi)* and *npp-22*$^{ndc1}$*(RNAi)* treated worms expressing markers for chromatin (magenta) and the plasma membrane (green). Images from the turn and central gonad region are stitched together and prominent signature defects for indicated RNAi conditions are highlighted. These include: large apoptotic regions (asterisks), unusual moon-shaped nuclei within the rachis region (box, zoom region), and prevalent compartmental bridging across the rachis (arrow), as indicated. Scale bar is 10 mm. D. CRISPR-Cas9 genome editing strategy used to generate the *npp-28Δ* deletion allele and the endogenous NPP-28ˆGFP strain. E. Immunoblot of lysates from N2, NPP-28ˆGFP, and *npp-28Δ* animals probed with anti-NPP-28 antibodies. α-tubulin served as the loading control. Red asterisk indicates non-specific bands. F. Brood size and embryonic lethality (bottom) of control and *npp-28Δ* animals. Each dot represents a single animal. N indicates number of animals scored and n is the total number of progeny. *** indicates p = 0.0005 and n.s is non-significant p = 0.5830 using Welch's t-tests on normal distributions as determined by the Shapiro-Wilke test. G. Representative spinning-disk confocal images of mCh::H2B embryos in control and *npp-28Δ* backgrounds during mitotic prophase. Time is shown relative to nuclear permeabilization in seconds. Insets highlight chromosome association with the nuclear periphery. Schematics illustrate chromosome tethering defects. H. Schematic illustrating quantification of chromosome positioning. Plot shows the peripheral association factor (PAF), calculated as the ratio of mean chromatin signal at the nuclear periphery (NP) to that at the nuclear center (NC), relative to nuclear permeabilization. Shaded regions represent SD. I. Quantification of embryos exhibiting nuclear envelope breakdown (NEBD) delays at anaphase onset the indicated strain. *n* indicates the number of embryos analyzed. Scale bars, 10 µm. Insets 5 µm.

### Puccini et al. Figure S2

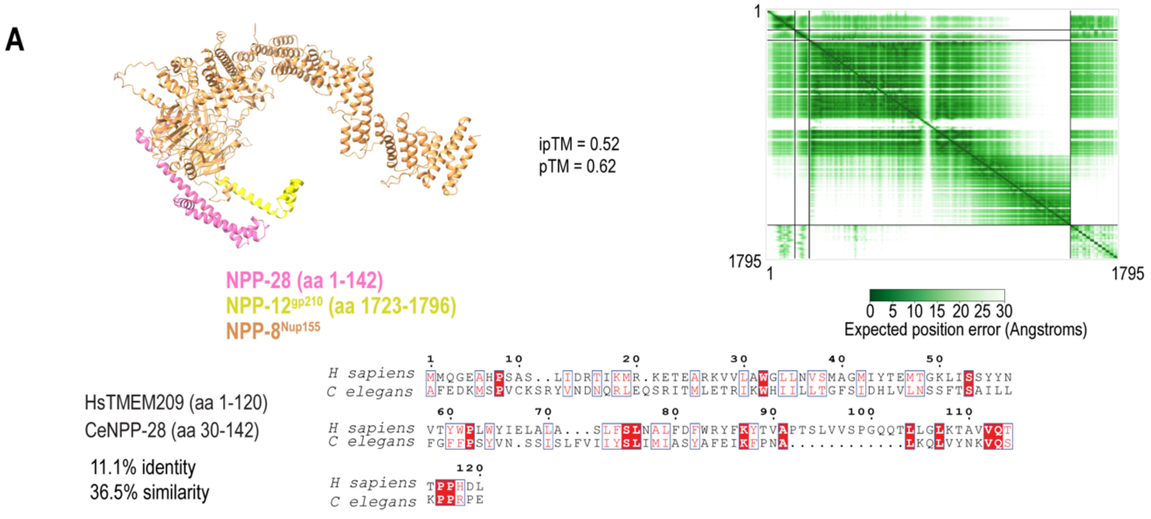


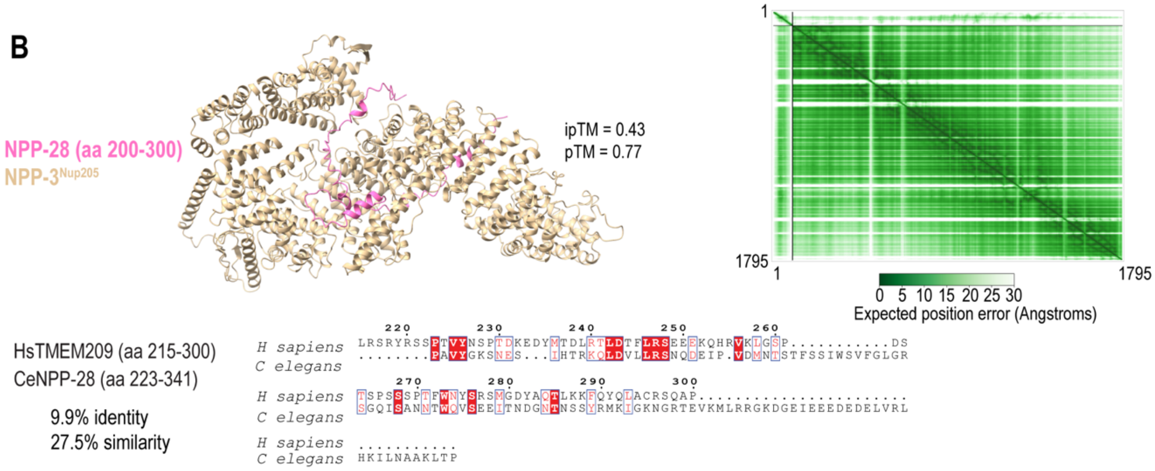


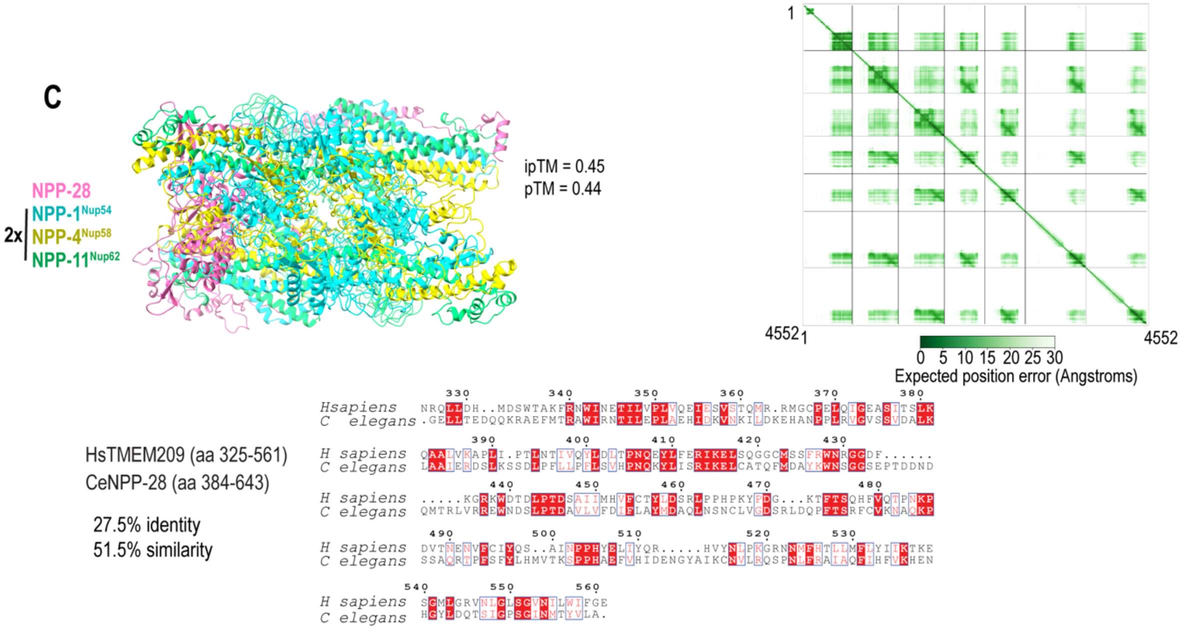

**Figure S2. Predicted interactions between NPP-28 and nuclear pore complex components.**

**A.** AlphaFold 3-predicted interaction between the N-terminal region of NPP-28 and NPP-12/gp210. Predicted aligned error (PAE) plot and sequence alignment between the corresponding regions of *C. elegans* NPP-28 and human TMEM209 are shown. **B.** AlphaFold 3-predicted interaction between the middle region of NPP-28 and NPP-3/Nup205. Predicted aligned error (PAE) plot and sequence alignment between the corresponding regions of *C. elegans* NPP-28 and human TMEM209 are shown. **C.** Alpha Fold 3-predicted interaction NPP-28 and 2 copies of the central channel complex components NPP-1/NPP-4/NPP-11 (Nup54/58/62). Predicted aligned error (PAE) plot and sequence alignment between the corresponding regions of *C. elegans* NPP-28 and human TMEM209 are shown. ipTM and pTM scores from AlphaFold 3 are indicated for each predicted complex.

Puccini et al. Figure S3

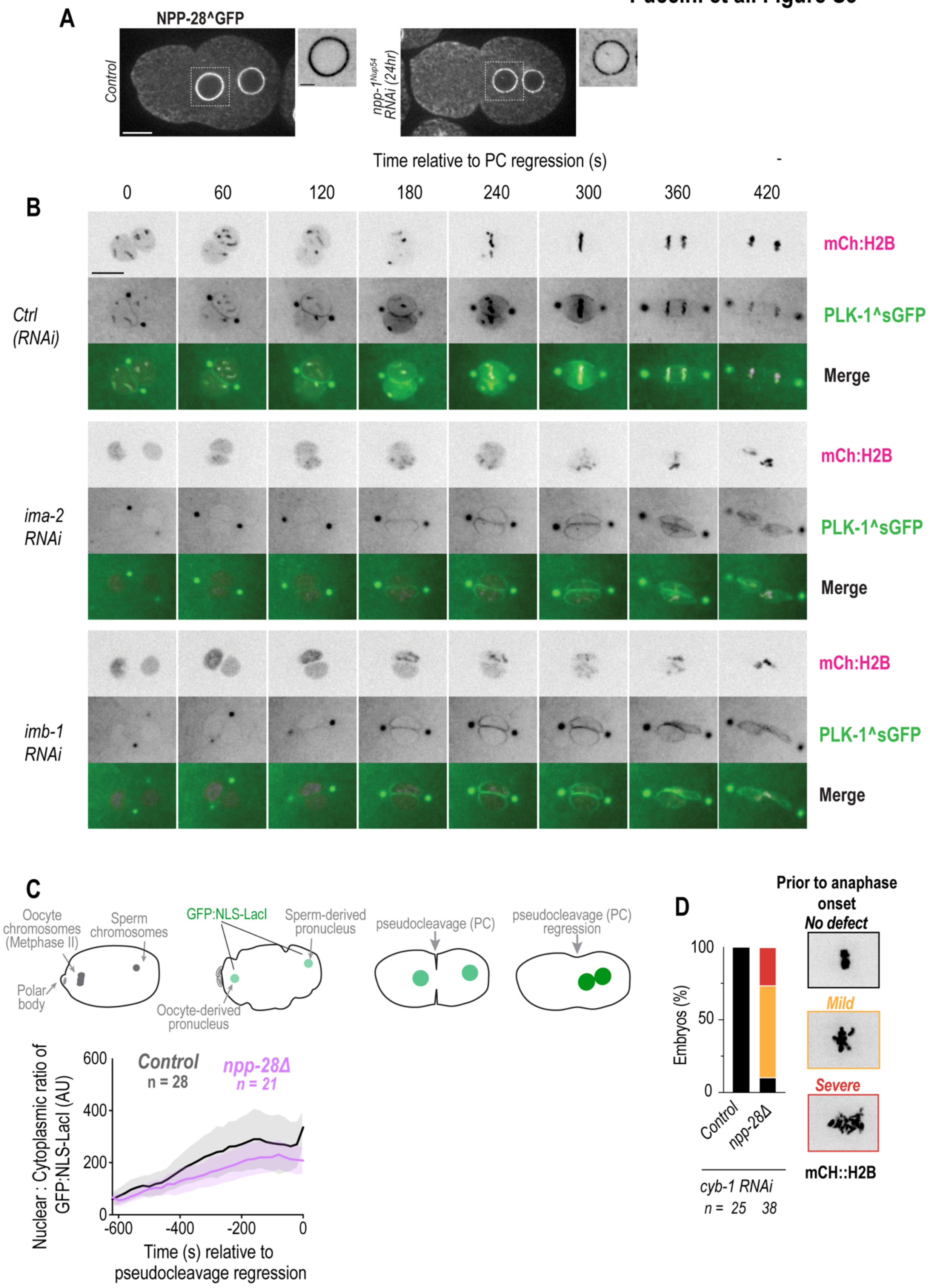

**Figure S3. NPP-28 regulates PLK-1 distribution independently of global defects in nuclear import.** A. Representative spinning disk confocal images of NPP-28^GFP in control and *npp-1(RNAi)* depleted early embryos at a time point prior to pseudocleavage regression. Zoom of female pronucleus shows NPP-28 distribution at the NE. B. Representative spinning-disk confocal images of PLK-1^sGFP and mCh::H2B embryos under the indicated RNAi conditions. Time shown is relative to pseudocleavage regression in seconds. C. (Top) Schematic illustrating quantification of the nucleoplasmic-to-cytoplasmic (N:C) ratio of GFP::NLS-LacI from fertilization to pseudocleavage regression. Plot shows the N:C ratio relative to pseudocleavage regression (PCR) in control and *npp-28Δ* embryos. Shaded regions represent SD. D. Plot of chromosome congression phenotypes in indicated conditions. Categorization of no defect, mild and severe are depicted in zooms of max projection of mCherry::Histone2B. Scale bars, 10 µm. *n* indicates the number of embryos analyzed.

**Puccini et al. Figure S4**

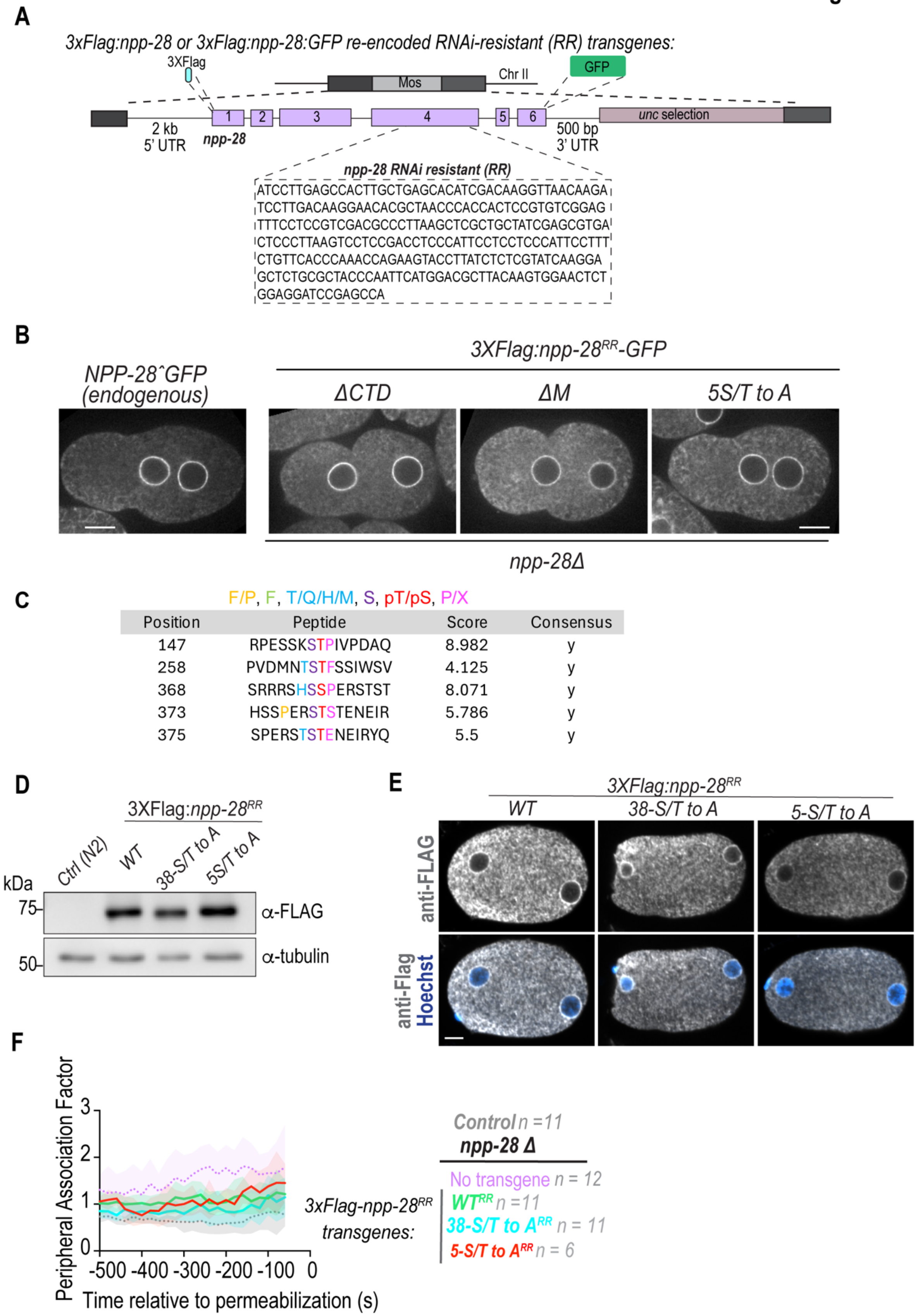

**Figure S4. Characterization of NPP-28 mutant transgenes used to assess PLK-1 docking and phosphorylation site function.** A. Schematic of the RNAi-resistant (RR) 3xFLAG::*npp-28* and 3xFLAG::*npp-28*::GFP transgenes. Re-encoded sequence used to confer RNAi resistance is indicated. B. Representative live cell spinning confocal images of endogenously tagged NPP-28^GFP and indicated 3xFLAG::NPP-28::GFP transgenes expressed in the *npp-28Δ* background. Scale bars, 10 µm. C. GPS-Polo 1.0 Predicted PLK-1 phospho-binding sites on NPP-28. Residues color coded according to PLK-1 PBD docking motif. Also in bottom of Table 1. D. Immunoblot of lysates from animals expressing the indicated 3xFLAG::NPP-28 transgenes probed with anti-FLAG antibodies. α-tubulin served as a loading control. E. Immunofluorescence images of embryos expressing the indicated 3xFLAG::NPP-28 RNAi-resistant (RR) transgenes stained with anti-FLAG antibodies. DNA was visualized with Hoechst. F. Plot shows the peripheral association factor (PAF), calculated as the ratio of mean chromatin signal at the nuclear periphery (NP) to that at the nuclear center (NC), relative to nuclear permeabilization, in indicated conditions (see also Supplemental Fig. S1F). Control and *npp-28*Δ conditions duplicated from Fig. S1F for reference. Shaded regions represent SD.

**Puccini et al. Figure S5**

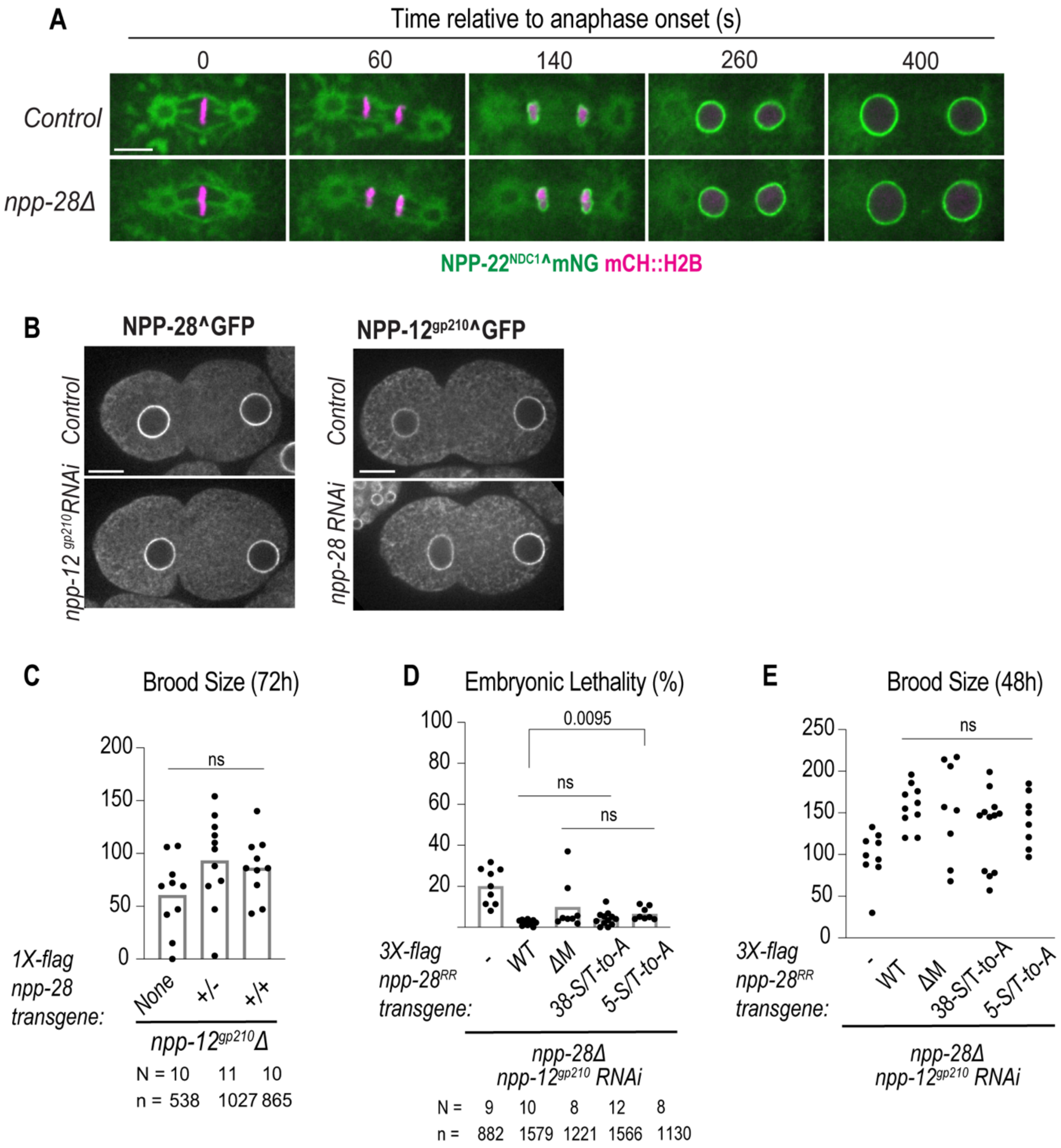

**Figure S5. Genetic interactions between *npp-28* and *npp-12* during embryogenesis and germline development. A.** Representative spinning-disk confocal images of NPP-22$^{NDC1}$^mNG and mCh::H2B embryos in control and *npp-28Δ* backgrounds. Time is shown relative to anaphase onset in seconds. **B.** Spinning disk confocal images of GFP tagged *npp-28* or *npp-12* at the endogenous locus under indicated conditions in early embryos prior to pseudocleavage regression (PCR). **C.** Brood size of *npp-12Δ* animals 72 h post L4 stage carrying zero, one, or two copies of indicated *npp-28* transgene in the endogenous *npp-28* background. Each dot represents a single animal. p values calculated with Brown-Forsythe and Welch ANOVA test with normal distributions, as calculated by a Shapiro-Wilk test. **D.** Embryonic lethality of *npp-28Δ; npp-12* (RNAi) embryos expressing the indicated 1xFlag::*npp-28* transgenes. Each dot represents an independent experiment. p values calculated using the Kruskal-Wallis test. **E.** Brood size of *npp-28Δ; npp-12* (RNAi) animals expressing the indicated 1xFlag::NPP-28 transgenes. Each dot represents a single animal. Each dot represents a single animal.  Scale bar, 10 µm. N indicates number of worms analyzed, *n* indicates the number of embryos. p values calculated with Brown-Forsythe and Welch ANOVA test with normal distributions, as calculated by a Shapiro-Wilk test.

**Supplemental Movie Legends**

**Movie 1. Similar nuclear envelope dynamics of NPP-28 and NPP-11$^{Nup62}$ with distinct ER and nuclear pools during the first mitotic division**, related to Figure 1. Spinning disk confocal microscopy time series of nuclear disassembly and assembly during the first cell division of *C. elegans* embryos. Endogenous NPP-28^GFP (green) and NPP-11$^{Nup62}$^mScarlet (magenta) are labeled. Images were acquired every 20 s – playback rate is 80X real time. Scale bar, 5 µm.

**Movie 2. Loss of *npp-28* results in delayed nuclear envelope disassembly and defects in assembly of GFP^NPP-19$^{Nup53}$**, related to Figure 2C. Spinning disk confocal microscopy time series of endogenous GFP^NPP-19$^{Nup53}$ nuclear disassembly and assembly during the first cell division of *C. elegans* embryos. Top, control; bottom; *npp-28Δ*. Images were acquired every 20 s and aligned to the onset of furrow ingression – playback rate is 80X real time. Scale bar, 5 µm.

**Movie 3. Loss of *npp-28* results in lack of nuclear envelope association of PLK-1^sGFP concomitant with an increased nuclear accumulation**, related to Figure 3. Spinning disk confocal time series of a single slice of early *C. elegans* embryos expressing endogenous PLK-1^sGFP. Images were acquired every 20 s and aligned to nuclear permeabilization, as determined by loss of the diffuse pool of mCh:H2B (not shown) – playback rate is 80X real time. Scale bar, 5 µm.

**Movie 4. Defective membrane clearance in *npp-12$^{gp210}$Δ* embryos is exacerbated by loss of *npp-28***, related to Figure 5. Spinning disk confocal time series of early *C. elegans* embryos expressing endogenous NPP-22$^{NDC1}$^mNG and mCh::H2B. Images were acquired every 20 s and aligned to anaphase onset – playback rate is 80X real time. Scale bar, 5 µm.

**Supplemental Tables**

**Table S1.** Excel workbook contains tables relevant to the nuclear screen, including: (1) top and reciprocal phenotypic matches for all genes, based on CSI scores, (2) manual scoring data for all genes- 153 parameters scored for each gene in a binary fashion, (3) descriptions of each scored parameter, and (4) forward and reverse oligo sequences used for the production of dsRNA.

**Table S2.** *C. elegans* strains used in this study

| ***C. elegans* strain name** | **Genotype** | **Source** |
|---|---|---|
| N2 | (Bristol): Wild-type (ancestral) | Caenorhabditis Genetics Center (CGC) |
| SBW479 | B0464.6(sbw23) III | This study |
| SBW791 | npp-11(ax4547[npp-11::wrmScarlet]) I; B0464.6(sbw46[B0464.6::GFP]) III | This study |
| OD95 | unc-119(ed3) III; ltIs37 [pAA64; pie-1/mCHERRY::his-58; unc119 (+)] IV; ltIs38 [pAA1; pie-1/GFP::PH(PLC1delta1); unc-119 (+)] | (De La Cruz Ruiz et al., 2022; Essex et al., 2009) |
| SBW313 | unc-119(ed3) III; ltIs37 [pAA64; pie-1p::mCherry::his-58 + unc-119 (+)] IV; ndc-1(sbw15[ndc1::mNG]) V | (Barger et al., 2026) |
| SBW488 | B0464.6(sbw23) III; unc-119(ed3) III; ltIs37 [pAA64; pie-1p::mCherry::his-58 + unc-119 (+)] IV; ndc-1(sbw15[ndc1::mNG]) V | This study |
| SBW575 | B0464.6 (sbw23) III - 6X outcrossed, his-72(cp76[mNG^3xFlag::his-72]) III | This study |
| LP312 | his-72(cp76[mNG^3xFlag::his-72]) III | (Dickinson et al., 2015) |
| WLP996 | jfSi68[lmn-1(4 kb 5'UTR)::mCherry::lmn-1gDNA exon four recoded::3'UTR lmn-1; cb-unc-119(+)]II; ojIs1[pie-1p::GFP::tbb-2 + unc-119(+)] IV | (Velez-Aguilera et al., 2022) |
| SBW768 | B0464.6(sbw46[B0464.6::GFP]) III | This study |
| OD270 | unc-119(ed3) III; ojls23[SP12::GFP unc-119(+)]; ltIs37 [pAA64; pie-1/mCHERRY::his-58; unc-119 (+)] | (Bahmanyar et al., 2014) |

| | | |
|---|---|---|
| SBW803 | B0464.6 (sbw23) III - 6X outcrossed; unc-119(ed3) III; ojls23[SP12::GFP unc-119(+)]; ltIs37 [pAA64; pie-1/mCHERRY::his-58; unc-119 (+)] IV | This study |
| JH4201 | npp-11(ax4547[npp-11::wrmScarlet]) I. | (Thomas et al., 2023) |
| SBW766 | npp-11(ax4547[npp-11::wrmScarlet]) I; B0464.6 (sbw23) III | This study |
| SBW814 | npp-19(bq29[G>F>P::npp-19]) II; unc-119(ed3) III; ltIs37 [pAA64; pie-1p::mCherry::his-58 + unc-119 (+)] IV | This study |
| SBW815 | npp-19(bq29[G>F>P::npp-19]) II; B0464.6(sbw23) III; unc-119(ed3) III; ltIs37 [pAA64; pie-1p::mCherry::his-58 + unc-119 (+)] IV | This study |
| SBW605 | mel-28(bq5[GFP::mel-28]) III; unc-119(ed3) III; ltIs37 [pAA64; pie-1p::mCherry::his-58 + unc-119 (+)] IV | This study |
| OD1496 | unc-119(ed3)III; ltSi464[pNH103; mex-5p::npp-6::GFP::tbb-2:3'UTR; cb-unc-119(+)]I; unc-119(ed3) III; ltIs37 [pAA64; pie-1p::mCherry::his-58 + unc-119 (+)] IV | (Hattersley et al., 2016) |
| JH3867 | bqSi189 [lmn-1p::mCherry::his-58::pie-1 3'utr] II; npp-10(ax4538[mNeonGreen::npp-10]) III | (Thomas et al., 2023) |
| SBW573 | plk-1(lt18[plk-1::sGFP]::loxp) III; unc-119(ed3) III; ltIs37 [pAA64; pie-1p::mCherry::his-58 + unc-119 (+)] IV | This study |
| SBW47 | unc-119(ed3) III; ltIs37 [pAA64; pie-1p::mCherry::his-58 + unc-119 (+)] IV; ltIs75 [(pSK5) pie-1p::GFP::TEV-STag::LacI + unc-119(+)] | (Penfield et al., 2020) |
| SBW811 | B0464.6(sbw23) III; unc-119(ed3) III; ltIs37 [pAA64; pie-1p::mCherry::his-58 + unc-119 (+)] IV; ltIs75 [(pSK5) pie-1p::GFP::TEV-STag::LacI + unc-119(+)] | This study |
| OD2591 | ltSi814[pPLG047; Pfzy-1::gfp::fzy-1::fzy-1 3'UTR; cb-unc-119(+)]I; unc-119(ed3)III?;; ltIs37 [pAA64; pie-1/mCHERRY::his-58; unc-119 (+)] IV | (Kim et al., 2017) |
| SBW821 | unc-119(ed3)III; sbwSi27 [pSB1065; 1xFlag::B0464.6] cb-unc-119(+) II; B0464.6(sbw23) III; unc-119(ed3) III; ltIs37 [pAA64; pie-1p::mCherry::his-58 + unc-119 (+)] IV; ndc-1(sbw15[ndc1::mNG]) V | This study |

| | | |
|---|---|---|
| SBW808 | unc-119(ed3)III; sbwSi28 [pSB1066;1xFlag::B0464.6ΔM]] cb-unc-119(+) II; B0464.6(sbw23) III; unc-119(ed3) III; ltIs37 [pAA64; pie-1p::mCherry::his-58 + unc-119 (+)] IV; ndc-1(sbw15[ndc1::mNG]) V | This study |
| SBW764 | unc-119(ed3)III; sbwSi29 [pSB1067;1xFlag::B0464.6ΔCTD] cb-unc-119(+) II; B0464.6(sbw23) III; unc-119(ed3) III; ltIs37 [pAA64; pie-1p::mCherry::his-58 + unc-119 (+)] IV; ndc-1(sbw15[ndc1::mNG]) V | This study |
| SBW789 | unc-119(ed3)III; sbwSi32 [pSB1068; B0464-5'UTR::3xFlag:B0464 (RNAi-resistant)::3'UTR-B0464] cb-unc-119(+) II | This study |
| SBW794 | unc-119(ed3)III; sbwSi33 [pSB1069; B0464-5'UTR::3xFlag:B0464-phospho-null (RNAi-resistant)::3'UTR-B0464] cb-unc-119(+) II | This study |
| SBW800 | unc-119(ed3)III; sbwSi35 [pSB1070; B0464-5'UTR::3xFlag:B0464-docking-mutant (RNAi-resistant)::3'UTR-B0464] cb-unc-119(+) II | This study |
| SBW792 | unc-119(ed3)III; sbwSi32 [pSB1068; B0464-5'UTR::3xFlag:B0464 (RNAi-resistant)::3'UTR-B0464] cb-unc-119(+) II; plk-1(lt18[plk-1::sGFP]::loxp) III; ltIs37 [pAA64; pie-1p::mCherry::his-58; unc-119 (+)] IV | This study |
| SBW797 | unc-119(ed3)III; sbwSi33 [pSB1069; B0464-5'UTR::3xFlag:B0464-phospho-null (RNAi-resistant)::3'UTR-B0464] cb-unc-119(+) II; plk-1(lt18[plk-1::sGFP]::loxp) III; ltIs37 [pAA64; pie-1p::mCherry::his-58; unc-119 (+)] IV | This study |
| SBW812 | unc-119(ed3)III; sbwSi35 [pSB1070; B0464-5'UTR::3xFlag:B0464-docking-mutant (RNAi-resistant)::3'UTR-B0464] cb-unc-119(+) II; plk-1(lt18[plk-1::sGFP]::loxp) III; ltIs37 [pAA64; pie-1p::mCherry::his-58 + unc-119 (+)] IV | This study |
| SBW829 | unc-119(ed3)III; sbwSi38 [pSB1071; B0464-5'UTR::3xFlag:B0464-deltaM (RNAi-resistant)::3'UTR-B0464] cb-unc-119(+) II; plk-1(lt18[plk-1::sGFP]::loxp) III; ltIs37 [pAA64; pie-1/mCHERRY::his-58; unc-119 (+)] IV | This study |
| SBW825 | unc-119(ed3)III; sbwSi32 [pSB1068; B0464-5'UTR::3xFlag:B0464 (RNAi-resistant)::3'UTR-B0464] cb-unc- | This study |

| | | |
|---|---|---|
| | 119(+) II; B0464.6(sbw23) III unc-119(ed3) III; ltIs37 [pAA64; pie-1p::mCherry::his-58 + unc-119 (+)] IV; ndc-1(sbw15[ndc1::mNG]) V | |
| SBW826 | unc-119(ed3)III; sbwSi33 [pSB1069; B0464-5'UTR::3xFlag:B0464-phospho-null (RNAi-resistant)::3'UTR-B0464] cb-unc-119(+) II; B0464.6(sbw23) III unc-119(ed3) III; ltIs37 [pAA64; pie-1p::mCherry::his-58 + unc-119 (+)] IV; ndc-1(sbw15[ndc1::mNG]) V | This study |
| SBW827 | unc-119(ed3)III; sbwSi35 [pSB1070; B0464-5'UTR::3xFlag:B0464-docking-mutant (RNAi-resistant)::3'UTR-B0464] cb-unc-119(+) II; B0464.6(sbw23) III; unc-119(ed3) III; ltIs37 [pAA64; pie-1p::mCherry::his-58 + unc-119 (+)] IV; ndc-1(sbw15[ndc1::mNG]) V | This study |
| SBW836 | unc-119(ed3)III; sbwSi39 [pSB1072; B0464-5'UTR::3xFlag:B0464-deltaM (RNAi-resistant)::GFP::3'UTR-B0464] cb-unc-119(+) II | This study |
| SBW838 | unc-119(ed3)III; sbwSi40 [pSB1073;B0464-5'UTR::3xFlag:B0464-deltaCTD::GFP::3'UTR-B0464] cb-unc-119(+) II | This study |
| SBW842 | unc-119(ed3)III; sbwSi41 [pSB1074;B0464-5'UTR::3xFlag:B0464-docking mutant::GFP::3'UTR-B0464] cb-unc-119(+) II | This study |
| SBW806 | npp-12(tm2320) I; unc-119(ed3)III; sbwSi27 [pSB1065; 1xFlag::B0464.6] cb-unc-119(+) II | This study |
| SBW759 | npp-12(tm2320) I; unc-119(ed3) III; ltIs37 [pAA64; pie-1p::mCherry::his-58 + unc-119 (+)] IV; ndc-1(sbw15[ndc1::mNG]) V | This study |
| DG4460 | npp-12(tn1780[npp-12:gfp:tev:3xFlag] I | (Huelgas-Morales et al., 2020) |

**Table S3.** Oligonucleotides used in this study.

| **Oligonucleotides for dsRNA production** | | | |
|---|---|---|---|
| Gene | Oligonucleotide 1 | Oligonucleotide 1 | Template |
| *B0464.6* | TAATACGACTCACTATAGGA CGTGGCAAAGATGGAGAGA | TAATACGACTCACTATAGGT CTGAACCACCCGAATTCCA | gDNA |

| | | | |
|---|---|---|---|
| *B0464.6 targeting re-encoded region* | TAATACGACTCACTATAGGT GCAAATCCTCCACTTCGTG | TAATACGACTCACTATAGGT CTGAACCACCCGAATTCCA | gDNA |
| *npp-1* | TAATACGACTCACTATAGGA TTTCAGGAATCGATGAGCG | TAATACGACTCACTATAGGA CTCGTAAAGGCCAGAGCAA | gDNA |
| *npp-12* | TAATACGACTCACTATAGGGc gacttctggaaaggtgag | TAATACGACTCACTATAGGT TGAGTTGGCACGAAGAGTG | gDNA |
| *ima-2* | TAATACGACTCACTATAGGC TGGAAATGACTCGCTGACA | TAATACGACTCACTATAGGT GAGTGATGATGCGGTAAGC | gDNA |
| *imb-1* | TAATACGACTCACTATAGGA GCTCCGAGCACGTATCAAT | TAATACGACTCACTATAGGC GATGGAGACGATGATGATG | gDNA |
| *ima-3* | TAATACGACTCACTATAGGC TTGATTGGCAGTGGGATTT | TAATACGACTCACTATAGGA CGTGTCCAAGGAGTGGAAC | cDNA |
| *cyb-1* | TAATACGACTCACTATAGGA TGCTAAGAGCCACCAACAAC | TAATACGACTCACTATAGGC GACTGGAGCTGGATTGGA | cDNA |

**Table S4.** Recombinant DNA or ssODN used in this study.

| Identifier | Recombinant DNA | Source |
|---|---|---|
| *B0464.6* crRNA-1 | AAGAGTGTCTAATTTATCGT**CGG** | IDT |
| *B0464.6* crRNA-2 | TATCTGGATCAGACCTCAAT**CGG** | IDT |

| | | |
|---|---|---|
| *B0464-GFP* repair template | gttgtgacatgtgacgtttccgattgaagatccggaagaatgtctaatatatcgtcggacatggtgatctgt<br>aaattaatatatttaaatatttaagacaattaaatagaaagctatatttattaaaattgagccaaaaaaataaaa<br>ttattaaaaatctccgaaatggcctagaaaatgctgacatgcgttttctaggccaacgcgctcgtagtacac<br>gaggacaaaaagattcgaaaatattttatcgaaaactccctcgaaaatgtttttctgtgatttttctcactaatt<br>tgttgttttaatataaatattaatctaatcgcaagctttacatcaactctagtgtgttacaggttcacaATGgt<br>ccttcttgagtttgtaacagctgctgggattacacatggcatggacgaactatacaaaagcccacaagcttt<br>gatgggcacaaatggaggagtcattgcggaacagtcaatggaaatcgaaacaaatgaaaacccagata<br>aggtagaggagcccgtcgtgagaaggaaacgagtaacccgcaggcgccacaggcggatacacagc<br>aagaataactgcttgacacctccaaacagcgatgacgaccctcaaatgagcacaccagatgatccagtc<br>atccacagtccaccatcaattggagcagctcctggtatgaatgggtatcatggatcaggagtgaaacttg<br>aagaaagttctggtgcttgtggctctccggatgatggcctgttagactccagcgaagagagtcgacgac<br>gtcagaaaacatgtagagtatgtggagatcatgcgacgggctataatttcaacgttatcacatgcgaatctt<br>gtaaagcattcttcaggagaaatgcgttacgtcccaaggagttcaaatgtccatattctgaagattgcgaa<br>attaattctgtaagcagacggttttgtcagaaatgtcgactgaggaaatgcttcacagttggtatgaaaaaa<br>gagtggattttaaacgaagaacaactgcggcggaggaaaaactctcgtttgaacaatacgggaacgtgt<br>aataaaagatcccagccagggaatcagcagtcaccacaaggtcctaatcaacagccacatttatcaccg<br>catcatccaggagttgcaatatatccaccacaaccacaacgcccactaacaatcaatccaatggacaatc<br>aaatgatgcatcatatgcaagctaacagaccaaatgccatgccacaacttatcagtccaccaggagcaca<br>accgtatccattgacatccccagtcggctctagtgcatctgactctccaccaaaccgaagtttaacaatgat<br>gcacaacggcgaaaaaagtccagatggctatgatccaaacattatggctcatcgtgccccgccgccatc<br>cttcaacaataggcctaaaatggatagtggacaggttgttctttccacggaagaatataaacaactcttgtc<br>tcgaattcgagccaatatgcgagaacacccgagaaaattcatcgatgatggtGGAGGAGGTG<br>GTGGAGGTATGACTGCGGATTCACTCCGCGAACGACTGAAAAG<br>T | Twist Bioscience |
| *B0464-phospho-null gene block* | ATGGATTACAAGGATCACGACGGAGACTATAAAGATCATGAT<br>ATCGACTACAAGGATGATGATGATAAGACTGCGGATGCTCTCC<br>GCGAACGACTGAAAAGTCAAAATGCAGACTATGCATTACTCAA<br>CACAGCTCAGGCAGCAAACCGGGCTGCATTTGAAGACAAAAT<br>GTCCCCAGTGTGCAAGAGCAGATATGTGAATGATAATCAGAGA<br>CTAGAACAAGCAAGAATAACAATGCTGGAAACCAGGATCAAA<br>TGGCACATCATTCTCTTAACTGGCTTCTCTATTGATgtaagagatgaaa<br>ctattaatttttattaaccagtttgtttcagCATTTGGTGCTCAATTCAGCTTTTACC<br>TCTGCAATTCTTCTTTTCGGATTCTTTCCCTCCTACGTAAATTCT<br>GCAATTTCACTATTCGTCATAATTTATTCATTGATTATGATTGC<br>ATCGTACGCGTTTGAGATCAAATTCCCGAATGCCCTGAAGCgtaa<br>gattttcagttattctaaaaaaacttattctatcgtttttcagAGCTGGTTTACAACAAAGTT<br>CAATCAAAACCACCACGTCCAGAATCAGCTAAAAGTACACCA<br>ATAGTGCCCGATGCTCAGAATGTGCTCGACACAGCAGTACAAA<br>GCAATGACCTTGCTTGGGTTGATGCTCACCGTTTCGGTACTCCA<br>GCATTCAAATCAGCTCAACTTCAACAAGCACCTGCTCCAAACA<br>AGAAGGCACCTGCTTATGCACAAGTGATGTCAAATGTTGCTGT<br>CGCAGACACTGCTGGAATTCTGGAGGATGCAAAAGGGGGATG<br>GAAAAGCCCGGCAGTTTATGGAAAATCTAATGAAGCTATTCAT<br>ACAAGAAAACAGCTTGATGTGCTTCTCCGATCTAATCAGGATG<br>AGATTCCAGTCGATATGAATACTGCAACATTTTCTTCCATTTGG<br>AGTGTATTTGGTTTAGGACGCAGTGGTCAGATAAGTGCGAATA<br>ACGCTTGGCAGGTTGCAGAGGAAATCGCTAATGATGGGAATAC<br>TAACTCAGgtaagttgcatcttcggtccaccatttcaaaaaaaaattgtgaaattttcaagatttttcat<br>tacaaaatttggttggtttgggaaattttgagtccgtcactccaccaacaaactatgctttaaatcatcaaattt<br>gtattatttttcagCATACCGAATGAAAATTGGTAAAAATGGAAGAACG<br>GAAGTAAAAATGTTGAGACGTGGCAAAGATGGAGAGATCGAG | Twist Bioscience and this study |

| | | |
|---|---|---|
| | GAAGAGGATGAAGACGAACTTGTCCGTTTGCATAAAATACTGA<br>ATGCGGCGAAGTTAACTCCCGAAGGAAAGACTGGAATTCTGA<br>AACGAAGCAATGCTATCGATCGTGCCGGGATTCGTAGTCGGCG<br>TAGAGCACATGCTTCACCTGAACGAGCAACTAGTGCTGAAAAT<br>GAAATCAGATACCAAACTGGAGAATTATTAACGGAAGATCAG<br>CAGAAAAGAGCCGAGTTTATGACTAGAGCCTGGATTCGCAACG<br>CAATCCTTGAGCCACTTGCTGAGCACATCGACAAGGTTAACAA<br>GATCCTTGACAAGGAACACGCTAACCCACCACTCCGTGTCGGA<br>GTTTCCTCCGTCGACGCCCTTAAGCTCGCTGCTATCGAGCGTGA<br>CTCCCTTAAGTCCTCCGACCTCCCATTCCTCCTCCCATTCCTTTC<br>TGTTCACCCAAACCAGAAGTACCTTATCTCTCGTATCAAGGAG<br>CTCTGCGCTACCCAATTCATGGACGCTTACAAGTGGAACTCTG<br>GAGGATCCGAGCCAGCTGATGATAACGATCAAATGACACGTCT<br>TGTTCGCCGTGAATGGAATGATGCATTGCCAACTGATGCAGTT<br>CTCGTATTCGACATTTTTCTGGCTTATATGGATGCTCAACTCAA<br>TTCCAATTGTCTCGTTGGAGATAGTCGCCTTGACCAGCCATTCA<br>CCAGCCGATTCTGTGTCAAAAATGCTCAAAAACCATCATCTGC<br>TCAACGTACTCCATTCAGTTTCTATCTTCACATGGTTACAAAAT<br>CTCCTCCACACGCTGAATTTGTGCACATTGATGAAAATGGATA<br>TGCAATTAAGTGCAACGTTCTCCGTCAATCACCCAATCTCTTCC<br>GTGCTATCGCTCAATTTATTCATTTTGTGAAACATGAAAATCAT<br>GGATATCTGGATCAGGCTTCAATCGGACCATCAGGAATTAACA<br>TGGCATACGTCCTGGCG | |
| *B0464-docking mutant gene block* | TCACTCCGCGAACGACTGAAAAGTCAAAATGCAGACTATTCAT<br>TACTCAACACATCACAGGCATCAAACCGGTCAGCATTTGAAGA<br>CAAAATGTCCCCAGTGTGCAAGAGCAGATATGTGAATGATAAT<br>CAGAGACTAGAACAATCGAGAATAACAATGCTGGAAACCAGG<br>ATCAAATGGCACATCATTCTCTTAACTGGCTTCTCTATTGATgtaa<br>gagatgaaactattaatttttattaaccagtttgtttcagCATTTGGTGCTCAATTCATCG<br>TTTACCTCTGCAATTCTTCTTTTCGGATTCTTTCCCTCCTACGTA<br>AATTCTAGTATTTCACTATTCGTCATAATTTATTCATTGATTAT<br>GATTGCATCGTACGCGTTTGAGATCAAATTCCCGAATGCCCTG<br>AAGCgtaagattttcagttattctaaaaaaacttattctatcgtttttcagAGCTGGTTTACAA<br>CAAAGTTCAATCAAAACCACCACGTCCAGAATCATCAAAAAGT<br>GCTCCAATAGTGCCCGATGCTCAGAATGTGCTCGACACAAGTG<br>TACAAAGCAATGACCTTTCTTGGGTTGATGCTCACCGTTTCGGT<br>ACTCCATCATTCAAATCAAGCCAACTTCAACAATCTCCTTCTCC<br>AAACAAGAAGTCACCTTCATATTCACAAGTGATGTCAAATGTT<br>TCAGTCTCAGACACTTCGGGAATTCTGGAGGATACCAAAGGGG<br>GATGGAAAAGCCCGGCAGTTTATGGAAAATCTAATGAATCGAT<br>TCATACAAGAAAACAGCTTGATGTGCTTCTCCGATCTAATCAG<br>GATGAGATTCCAGTCGATATGAATACTTCTGCTTTTTCTTCCAT<br>TTGGAGTGTATTTGGTTTAGGACGCAGTGGTCAGATAAGTGCG<br>AATAACACTTGGCAGGTTTCTGAGGAAATCACGAATGATGGGA<br>ATACTAACTCATgtaagttgcatcttcggtccaccatttcaaaaaaaaattgtgaaattttcaaga<br>tttttcattacaaaatttggttggtttgggaaattttgagtccgtcactccaccaacaaactatgctttaaatcat<br>caaatttgtattatttttcagCATACCGAATGAAAATTGGTAAAAATGGAAG<br>AACGGAAGTAAAAATGTTGAGACGTGGCAAAGATGGAGAGAT<br>CGAGGAAGAGGATGAAGACGAACTTGTCCGTTTGCATAAAAT<br>ACTGAATGCGGCGAAGTTAACTCCCGAAGGAAAGACTGGAAT<br>TCTGAAACGAAGCAATTCAATCGATCGTGCCGGGATTCGTAGT | Twist Bioscience and this study |

| | | |
|---|---|---|
| | CGGCGTAGATCTCATAGTGCTCCTGAACGATCTGCTAGTGCTG AAAATGAAATCAGATACCAAACTGGAGAATTATTAACGGAAG ATCAGCAGAAAAGAGCCGAGTTTATGACTAGAGCCTGGATTCG CAACACA | |
| pSB1065 | pCFJ151-B0464.6-5'UTR-1xFlag-B0464.6-B0464.6-3'UTR | This study |
| pSB1066 | pCFJ151-B0464.6-5'UTR-1xFlag-B0464.6ΔM-B0464.6-3'UTR | This study |
| pSB1067 | pCFJ151- B0464.6-5'UTR-1xFlag-B0464.6 ΔCTD-B0464.6-3'UTR | This study |
| pSB1068 | pCFJ151-B0464.6-5'UTR-3xFlag-B0464.6-re-encoded-B0464.6-3'UTR | This study |
| pSB1069 | pCFJ151-B0464.6-5'UTR-3xFlag-B0464.6-phospho-null-re-encoded-B0464.6-3'UTR | This study |
| pSB1070 | pCFJ151-B0464.6-5'UTR-3xFlag-B0464.6-docking mutant-re-encoded-B0464.6-3'UTR | This study |
| pSB1071 | PCFJ151-B0464-5'UTR::3xFlag:B0464-deltaM (RNAi-resistant)::3'UTR-B0464 | This study |
| pSB1072 | PCFJ151- B0464-5'UTR::3xFlag:B0464-deltaM (RNAi-resistant)::GFP::3'UTR-B0464 | This study |
| pSB1073 | PCFJ151-B0464-5'UTR::3xFlag:B0464-deltaCTD::GFP::3'UTR-B0464 | This study |
| pSB1074 | PCFJ151-B0464-5'UTR::3xFlag:B0464-docking mutant::GFP::3'UTR-B0464 | This study |
